# Scalable Apparatus to Measure Posture and Locomotion (SAMPL): a high-throughput solution to study unconstrained vertical behavior in small animals

**DOI:** 10.1101/2023.01.07.523102

**Authors:** Yunlu Zhu, Franziska Auer, Hannah Gelnaw, Samantha N. Davis, Kyla R. Hamling, Christina E. May, Hassan Ahamed, Niels Ringstad, Katherine I. Nagel, David Schoppik

## Abstract

Balance and movement are impaired in a wide variety of neurological disorders. Recent advances in behavioral monitoring provide unprecedented access to posture and loco-motor kinematics, but without the throughput and scalability necessary to screen candidate genes / potential therapeutics. We present a powerful solution: a Scalable Apparatus to Measure Posture and Locomotion (SAMPL). SAMPL includes extensible imaging hardware and low-cost open-source acquisition software with real-time processing. We first demonstrate that SAMPL’s hardware and acquisition software can acquire data from *D. melanogaster*, *C.elegans*, and *D. rerio* as they move vertically. Next, we leverage SAMPL’s throughput to rapidly (two weeks) gather a new zebrafish dataset. We use SAMPL’s analysis and visualization tools to replicate and extend our current understanding of how zebrafish balance as they navigate through a vertical environment. Next, we discover (1) that key kinematic parameters vary systematically with genetic background, and (2) that such background variation is small relative to the changes that accompany early development. Finally, we simulate SAMPL’s ability to resolve differences in posture or vertical navigation as a function of effect size and data gathered – key data for screens. Taken together, our apparatus, data, and analysis provide a powerful solution for laboratories using small animals to investigate balance and locomotor disorders at scale. More broadly, SAMPL is both an adaptable resource for laboratories looking process video-graphic measures of behavior in real-time, and an exemplar of how to scale hardware to enable the throughput necessary for screening.

## INTRODUCTION

Measuring posture and locomotion is key to understand nervous system function and evaluate potential treatments for disease – particularly neurological disorders ^1^. Behavioral screening is a fundamental part of both basic and translational approaches to disease ^2, 3^. For screens, measuring behavior from large numbers of animals is necessary to differentiate individual variation ^4^ from changes seen in disease models and/or improvement following treatment ^5, 6^. The demand for such high-throughput measurements comes at a cost: often, measurements that require high resolution – such as posture – are limited. Modern machine learning algorithms and inexpensive videographic / computing hardware have automated measurements of posture and kinematics ^7–9^ and illuminated our understanding of animal behavior ^10–12^. We sought to combine videographic analysis of posture and vertical locomotion with the scalability amenable to screening.

Over the past decade, we have studied posture and locomotion using the larval zebrafish as a model. Neural architecture is highly conserved across vertebrates, making larval zebrafish an excellent model to understand the underpinnings of locomotion ^13, 14^ and balance ^15^. For our studies, we developed a new apparatus/analysis pipeline to measure the statistics of posture in the pitch (nose-up/nose-down) axis and locomotion as larvae swam freely in depth. We discovered that larvae learn to time their movements to facilitate balance ^16^, that larvae modulate the kinematics of swimming to correct posture ^17^, and that larvae engage their pectoral fins to climb efficiently ^18^, and implicated different neuronal circuits in each of these behaviors. While informative, data collection was slow (months) on small numbers (<5) of apparatus. Increasing throughput remains a challenge common to laboratories that develop new tools to measure behavior.

To meet the needs of scalability, resolution, and extensibility we developed SAMPL: a low-cost, open-source solution that measures posture and vertical locomotion in real-time in small animals. Further, we provide a turn-key analysis pipeline to measure larval zebrafish balance behavior. We begin with a brief treatment of the hardware and software; a detailed design guide, assembly and operating instructions are included as supplemental appendices. Next, we use SAMPL to measure unconstrained vertical locomotion in two common invertebrate models: flies (*Drosophila melanogaster*), and worms (*Caenorhabditis elegans*), as well as a small model vertebrate, the larval zebrafish (*Danio rerio*). To illustrate SAMPL’s capabilities, we parameterize a new dataset focused on behaviors that larval zebrafish perform as they stabilize posture and navigate (i.e. climb/dive) in the water column. Our new dataset represents two weeks worth of data collection, and allowed us to detail variation in postural/locomotor behaviors. By measuring behavior across different genetic backgrounds and development, we report two new findings. First, variation in posture/locomotion is systematic across genotype and second, the scale of variation in behavior across development is much larger than background genetic variation. We use these new data to simulate the resolving power for each behavioral parameter as a function of data gathered – foundational information to rigorously assay the effects of candidate genes or small molecules on posture or locomotion. SAMPL thus offers a straight-forward way to gather data from small animals, and a turn-key solution to screen for balance and vertical locomotion in larval zebrafish. More broadly, SAMPL offers a template for laboratories looking to scale their own behavioral apparatus to achieve the throughput necessary for screens. SAMPL will thus facilitate reproducible studies of postural and locomotor behaviors in both health and disease, addressing unmet needs in treating neurological disorders, particularly with balance symptoms ^19^.

## RESULTS

### SAMPL hardware & software overview

To overcome measure posture with the throughput necessary for genetic and drug screens, we deployed SAMPL, a real-time videographic system (Figure 1A) that records small animal behavior in the vertical axis. Below we briefly describe the hardware and software that comprise SAMPL. SAMPL’s hardware consists of three simple modules: an infrared (IR) illumination module (Figure 1B), a camera-lens module (Figure 1C), and two clamps to hold fish chambers (Figure 1D). All three modules are mounted directly (Figure 1A) onto an aluminum breadboard (Figure S1) and a light-tight enclosure covers the entire apparatus to permit individual control of lighting (Figures 1F and 1G). Details of hardware and software design can be found in Appendices 1&2. A complete parts list is in Table 1, hardware assembly instructions in Appendix 3, and a stop-motion movie of assembly provided as Movie 1.

**Figure 1:**
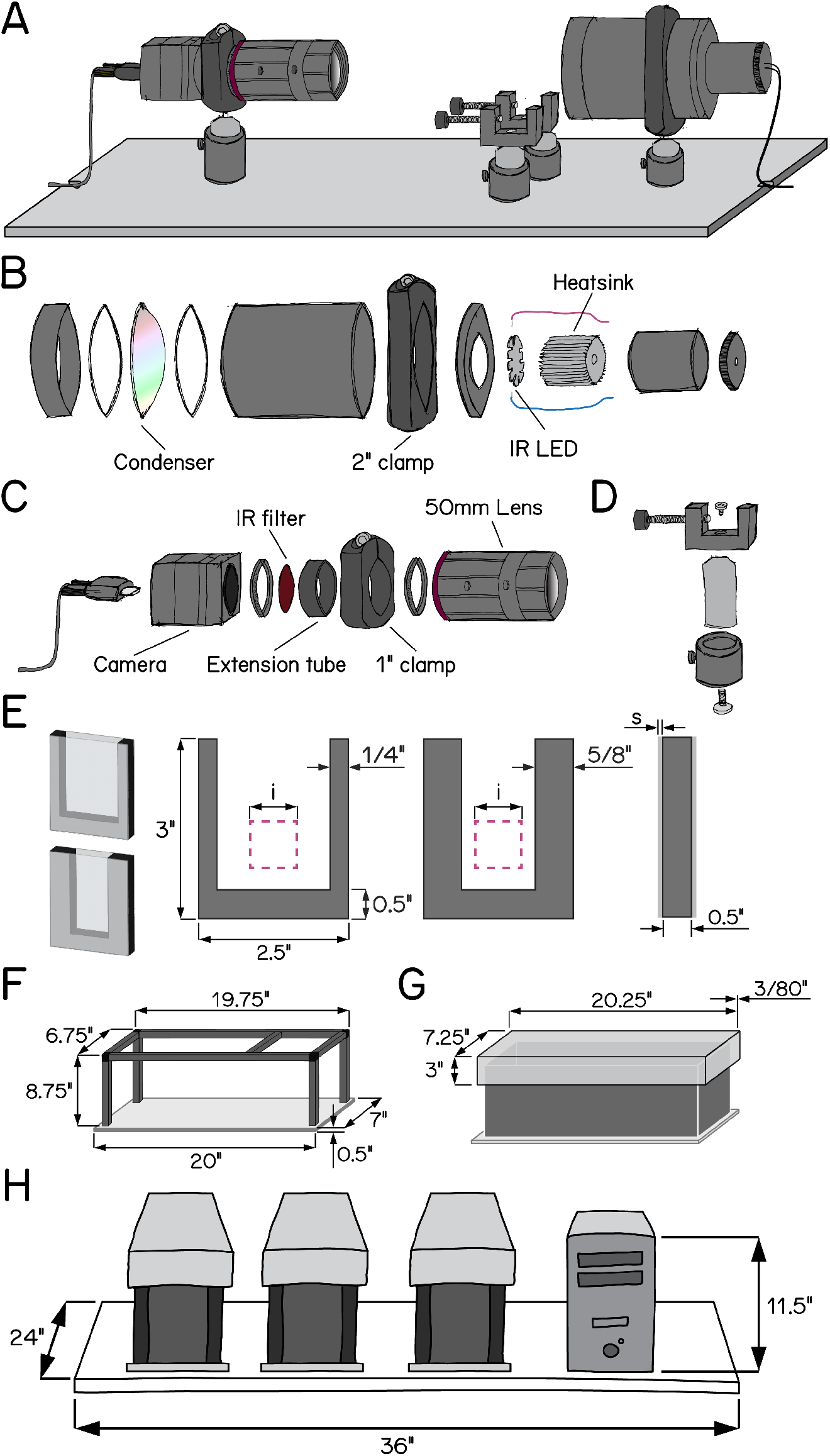
Schematic illustrations of SAMPL hardware design. (A) Overview of the apparatus without aluminum rails, side panels, and the top panel. Equipment modules mounted on the breadboard are, from left to right, IR camera and lens, chamber holders, and IR illumination module. (B) Exploded-view drawing of the IR illumination module. (C) Exploded-view drawing of the camera and lens module. (D) Exploded-view drawing of a chamber holder (E) Design of fish chambers. From left to right: 3D illustration of a standard chamber (upper) and a narrow chamber (lower); front view of the u-shaped acrylic middle piece for the chambers; side view of the chamber. Pink squares illustrate the recording field of view. i = 20 mm; s = 1.5 mm. (F) Dimensions of the apparatus frame and breadboard. (G) Design and dimensions of the apparatus lid. (H) Schematic illustration of a set of three SAMPL apparatus and a small-form-factor computer case on a 24”x36” shelf.

**Table 1:**
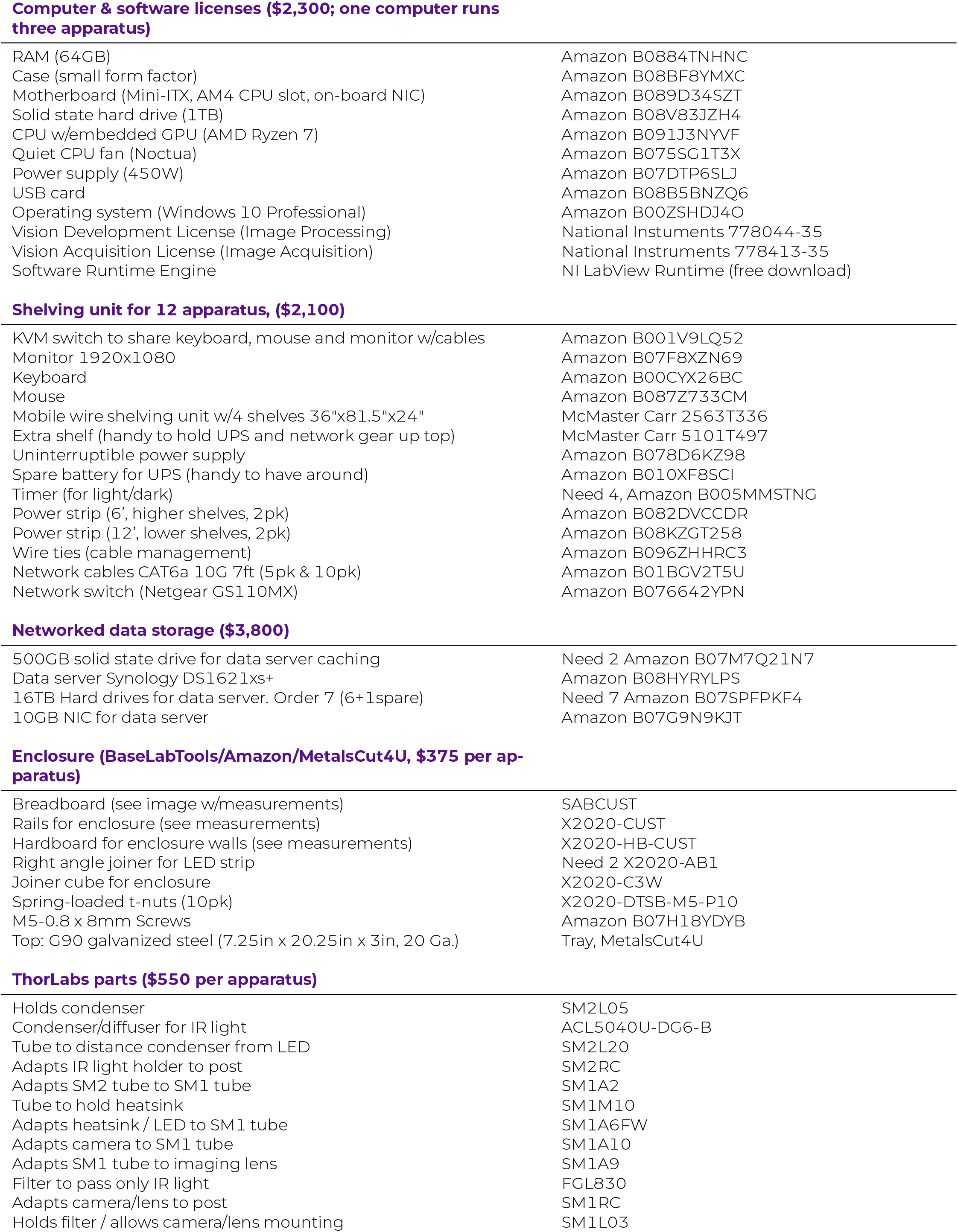
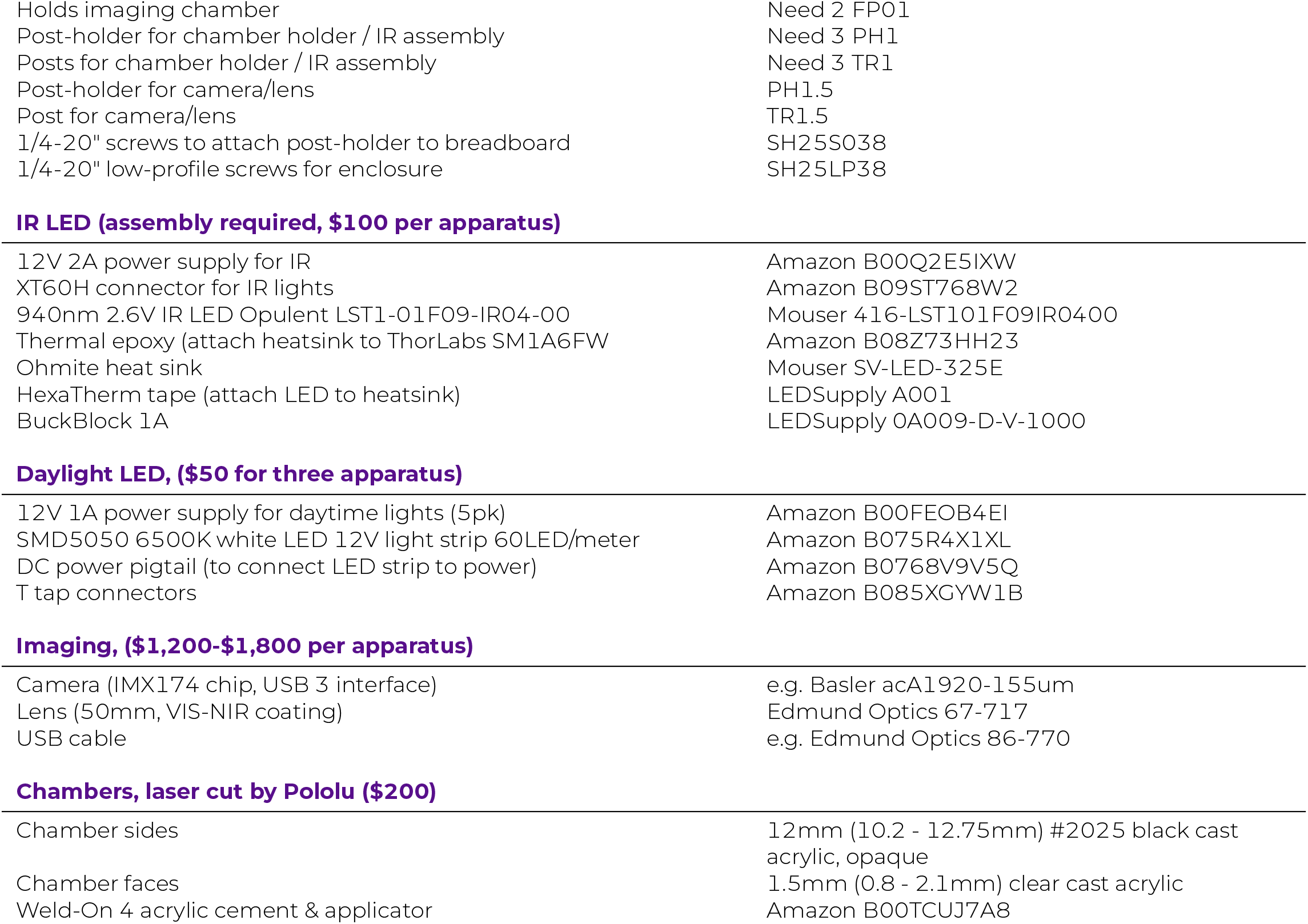
List of parts, prices per 12/2022

**Computer & software licenses (\$2,300; one computer runs three apparatus)**
|  |  |
| --- | --- |
| RAM (64GB) | Amazon B0884TNHNC |
| Case (small form factor) | Amazon B08BF8YMXC |
| Motherboard (Mini-ITX, AM4 CPU slot, on-board NIC) | Amazon B089D34SZT |
| Solid state hard drive (1TB) | Amazon B08V83JZH4 |
| CPU w/embedded GPU (AMD Ryzen 7) | Amazon B091J3NYVF |
| Quiet CPU fan (Noctua) | Amazon B075SG1T3X |
| Power supply (450W) | Amazon B07DTP6SLJ |
| USB card | Amazon B08B5BNZQ6 |
| Operating system (Windows 10 Professional) | Amazon B00ZSHDJ4O |
| Vision Development License (Image Processing) | National Instruments 778044-35 |
| Vision Acquisition License (Image Acquisition) | National Instruments 778413-35 |
| Software Runtime Engine | NI LabView Runtime (free download) |
| **Shelving unit for 12 apparatus, (\$2,100)** |  |
| KVM switch to share keyboard, mouse and monitor w/cables | Amazon B001V9LQ52 |
| Monitor 1920x1080 | Amazon B07F8XZN69 |
| Keyboard | Amazon B00CYX26BC |
| Mouse | Amazon B087Z733CM |
| Mobile wire shelving unit w/4 shelves 36"x81.5"x24" | McMaster Carr 2563T336 |
| Extra shelf (handy to hold UPS and network gear up top) | McMaster Carr 5101T497 |
| Uninterruptible power supply | Amazon B078D6KZ98 |
| Spare battery for UPS (handy to have around) | Amazon B010XF8SCI |
| Timer (for light/dark) | Need 4, Amazon B005MMSTNG |
| Power strip (6', higher shelves, 2pk) | Amazon B082DVCCDR |
| Power strip (12', lower shelves, 2pk) | Amazon B08KZGT258 |
| Wire ties (cable management) | Amazon B096ZHHR3C |
| Network cables CAT6a 10G 7ft (5pk & 10pk) | Amazon B01BGV2T5U |
| Network switch (Netgear GS110MX) | Amazon B076642YPN |
| **Networked data storage (\$3,800)** |  |
| 500GB solid state drive for data server caching | Need 2 Amazon B07M7Q21N7 |
| Data server Synology DS1621xs+ | Amazon B08HYRYLPS |
| 16TB Hard drives for data server. Order 7 (6+1spare) | Need 7 Amazon B07SPFPKF4 |
| 10GB NIC for data server | Amazon B07G9N9KJT |
| **Enclosure (BaseLabTools/Amazon/MetalsCut4U, \$375 per apparatus)** |  |
| Breadboard (see image w/measurements) | SABCUST |
| Rails for enclosure (see measurements) | X2020-CUST |
| Hardboard for enclosure walls (see measurements) | X2020-HB-CUST |
| Right angle joiner for LED strip | Need 2 X2020-AB1 |
| Joiner cube for enclosure | X2020-C3W |
| Spring-loaded t-nuts (10pk) | X2020-DTSB-M5-P10 |
| M5-0.8 x 8mm Screws | Amazon B07H18YDYB |
| Top: G90 galvanized steel (7.25in x 20.25in x 3in, 20 Ga.) | Tray, MetalsCut4U |
| **ThorLabs parts (\$550 per apparatus)** |  |
| Holds condenser | SM2L05 |
| Condenser/diffuser for IR light | ACL5040U-DG6-B |
| Tube to distance condenser from LED | SM2L20 |
| Adapts IR light holder to post | SM2RC |
| Adapts SM2 tube to SM1 tube | SM1A2 |
| Tube to hold heatsink | SM1M10 |
| Adapts heatsink / LED to SM1 tube | SM1A6FW |
| Adapts camera to SM1 tube | SM1A10 |
| Adapts SM1 tube to imaging lens | SM1A9 |
| Filter to pass only IR light | FGL830 |
| Adapts camera/lens to post | SM1RC |
| Holds filter / allows camera/lens mounting | SM1L03 |
| Holds imaging chamber | Need 2 FP01 |
| Post-holder for chamber holder / IR assembly | Need 3 PH1 |
| Posts for chamber holder / IR assembly | Need 3 TR1 |
| Post-holder for camera/lens | PH1.5 |
| Post for camera/lens | TR1.5 |
| 1/4-20" screws to attach post-holder to breadboard | SH25S038 |
| 1/4-20" low-profile screws for enclosure | SH25LP38 |
| IR LED (assembly required, \$100 per apparatus) |  |
| 12V 2A power supply for IR | Amazon B00Q2E5IXW |
| XT60H connector for IR lights | Amazon B09ST768W2 |
| 940nm 2.6V IR LED Opulent LST1-01F09-IR04-00 | Mouser 416-LST101F09IR0400 |
| Thermal epoxy (attach heatsink to ThorLabs SM1A6FW) | Amazon B08Z73HH23 |
| Ohmite heat sink | Mouser SV-LED-325E |
| HexaTherm tape (attach LED to heatsink) | LEDSupply A001 |
| BuckBlock 1A | LEDSupply 0A009-D-V-1000 |
| Daylight LED, (\$50 for three apparatus) |  |
| 12V 1A power supply for daytime lights (5pk) | Amazon B00FEOB4EI |
| SMD5050 6500K white LED 12V light strip 60LED/meter | Amazon B075R4X1XL |
| DC power pigtail (to connect LED strip to power) | Amazon B0768V9V5Q |
| T tap connectors | Amazon B085XGYW1B |
| Imaging, (\$1,200-\$1,800 per apparatus) |  |
| Camera (IMX174 chip, USB 3 interface) | e.g. Basler acA1920-155um |
| Lens (50mm, VIS-NIR coating) | Edmund Optics 67-717 |
| USB cable | e.g. Edmund Optics 86-770 |
| Chambers, laser cut by Pololu (\$200) |  |
| Chamber sides | 12mm (10.2 - 12.75mm) #2025 black cast acrylic, opaque |
| Chamber faces | 1.5mm (0.8 - 2.1mm) clear cast acrylic |
| Weld-On 4 acrylic cement & applicator | Amazon B00TCUJ7A8 |

The IR module illuminates the arena from behind. It is optimized to fulfill four criteria: (1) high image quality; (2) a large area for imaging; (3) imperceptible illumination; (4) ample heat dissipation. We used a 940 nm “star” style LED as our source of IR illumination and developed a simple illumination module to diffuse IR light across a 50mm circle (Figure 1B). For heat management, each LED was mounted to a small heat sink (Figure 1B). This setup allows us to power three illumination modules in series using a single LED driver.

The second module captures videographic data. It consists of a camera and lens optimized for speed, resolution, compactness, and affordability. The camera hardware satisfies the following demands: (1) large pixel size with low noise allowing for high dynamic range / signal-to-noise ratio; (2) sufficient resolution to resolve subtle changes to animal posture; (3) an interface with sufficient bandwidth for data transfer; (4) availability. The lens achieves (1) close focus; (2) sufficient depth-of-field to cover the entire depth of the imaging arena; (3) high image quality; (4) compact size; (5) high IR transmission rate; (6) ease of integrating an IR-pass filter. We adapted a 50 mm IR-optimized lens by placing a 0.3” extension tube between the lens and the camera to achieve higher magnification ratio with minimum working distance. The space between camera adapters and the extension tube allows us to fit a 25 mm IR-pass filter; the extension tube gives a mount point to connect the module to the base (Figure 1A). Using this camera-lens module, we image an area ∼400 mm^2^ (Figure 1E, pink square) at 166 Hz with 1200×1216 pixels at a focal distance of ∼24 cm.

The final module is a rectangular arena optimized for vertical locomotion (i.e. parallel to the focal plane). By design, the chamber size is larger than the imaging area, allowing stochastic sampling of freely behaving animals in a large enough arena. The bottom of the chamber is below the field of view so that animals sitting at the bottom will not be recorded. We assembled custom-fabricated chambers from laser-cut acrylic by cementing transparent front and back sides to a U-shaped piece that forms the narrower sides (Figure 1E). We designed two types of chambers with different inner widths to adapt to the needs of different experiments: a wider standard chamber optimized for larger groups of animals and a narrower chamber for 1-3 animals (Figure 1E). Chambers can be easily dropped into the holders (Figure 1D) from the top of the behavior box and secured in place for recording.

SAMPL includes a complete suite of open-source software for acquisition/real-time extraction of data (source and compiled executables provided). Acquisition consists of a graphical user interface, written in LabView that analyzes video in real-time to isolate an animal’s location and orientation, with the ability to save raw video for further off-line analysis. The real-time processing algorithm consists of: (1) background subtraction; (2) noise thresholding; (3) rejection of frames without an animal or with >1 animal in view; (4) size and intensity criteria to identify two distinct animal parts, usually the body and the head; (5) image processing to extract location and body orientation relative to the horizon. Data about location and orientation is saved to a text file, metadata about the experiment is saved to a separate text file, and optionally, video is saved as an AVI file.

SAMPL’s modules and software were designed to scale, minimizing footprint and experimenter time. We multiplex apparatus, providing three distinct compiled applications designed to run simultaneously on one computer to reduce cost/footprint. A set of three SAMPL apparatus and a computer case fit on one 24”x36” shelf (Figure 1H). One SAMPL “rack” consists of four such shelves (81.5” high) and costs ∼$40,000-$45,000 (December 2022, before volume discounts). In our laboratory, trained experimenters can load such a rack for a typical 48 hr experiment in 30 minutes. Taken together, SAMPL’s design is ideal to efficiently gather data describing posture and vertical locomotion.

### SAMPL validation: different small animals

SAMPL is well-suited to collect data from a wide range of small animals. We demonstrate the flexibility of SAMPL’s acquisition suite using three common model organisms. By changing SAMPL’s thresholds (Table 2), we could acquire data from three different organisms: *Drosophila melanogaster* climbing behavior (Figures 2A and 2B), continuous locomotion in *Caenorhabditis elegans* (Figures 2C and 2D), and swimming in *Danio rerio* (Figures 2E and 2F). We present raw video from the epochs in Figure 2 together with plots of real-time image processing (fly & worms, Movie 2; fish, Movie 3). These results demonstrate SAMPL’s excellent flexibility and robustness in real-time recording and analysis of vertical locomotion of small animals.

**Figure 2:**
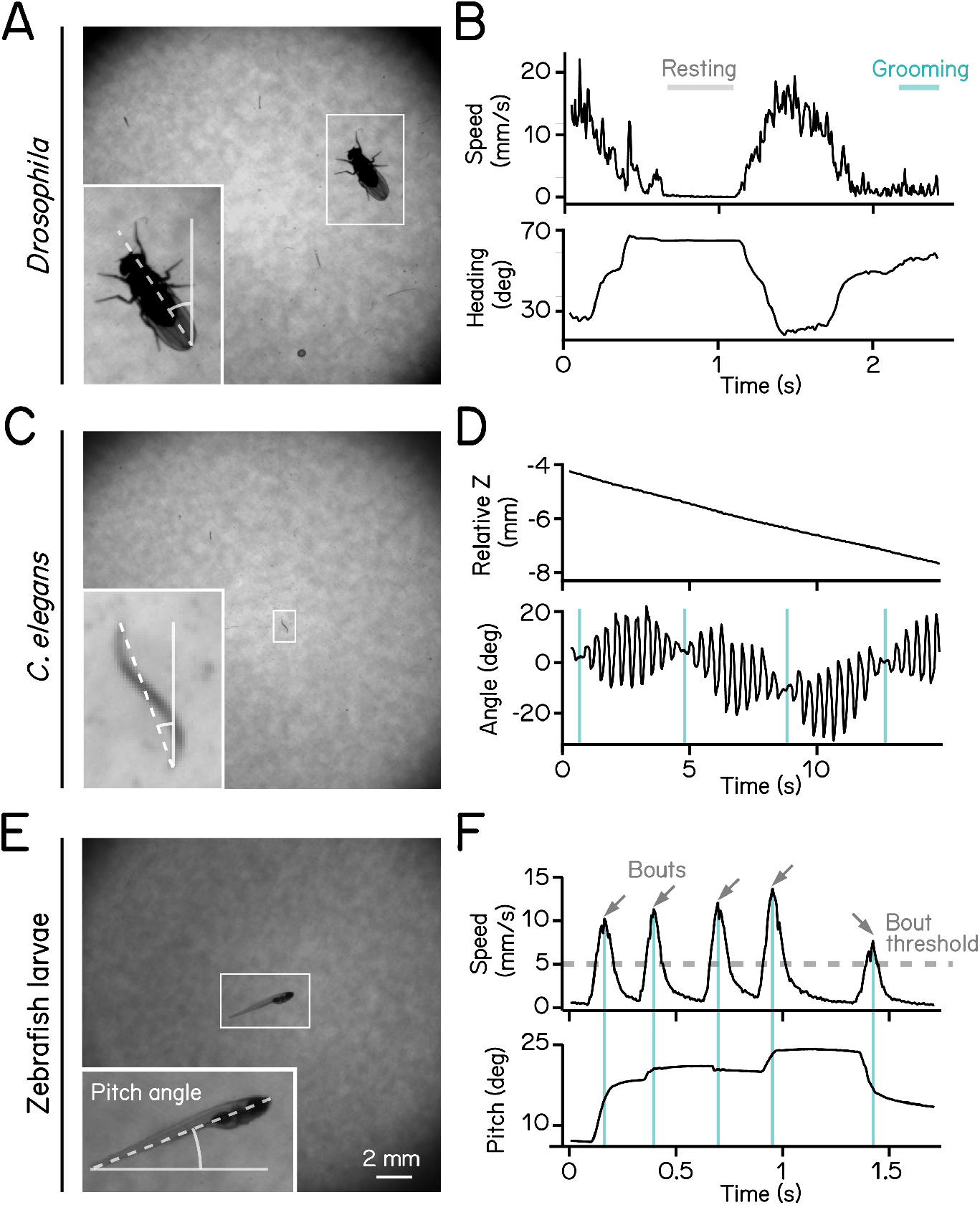
High-definition recording and measurement of animal locomotion using SAMPL. (A) Example of a recorded frame with a *Drosophila melanogaster* (white box) in the SAMPL apparatus. Dashed line indicates heading of the fly relative to vertical up (north). Imaging was performed at 166 Hz with 1200 1216 pixels. Same as follows. (B) Example of an epoch of a walking fly. Walking speed and heading are plotted as a function of time. Gray and cyan lines marks resting and grooming period, respectively (Movie 2). (C) Example of a recorded frame with a *Caenorhabditis elegans* (white box) in the SAMPL apparatus. Dashed line indicates approximated angle of the worm relative to vertical. (D) Example of an epoch of a swimming worm. Z position and approximated angle are plotted as a function of time. Cyan vertical lines label the time when the plane of movement is perpendicular to the imaging plane (Movie 2). (E) Example of a recorded frame with a 7 dpf *Danio rerio* larva (white box) in the SAMPL apparatus. Pitch angle is determined as the angle of the trunk of the fish (dashed line) relative to horizontal. Positive pitch indicates nose-up posture whereas negative pitch represents nose-down posture. (F) Example of an epoch containing multiple swim bouts (arrows). Swim speed and pitch angles are plotted as a function of time. Dashed line marks the 5 mm/s threshold for bout detection. Cyan vertical lines label time of the peak speed for each bout (Movie 3).

**Table 2:** Recording parameters for different organisms

| | Zebrafish ≤ 12 dpf | Zebrafish $>$ 12 dpf | *Drosophila* | *C. elegans* |
| --- | --- | --- | --- | --- |
| Body low | 14 | 14 | 100 | 20 |
| Body high | 255 | 255 | 255 | 255 |
| Head low | 45 | 45 | 30 | 21 |
| Head high | 255 | 255 | 255 | 255 |
| Initial cut low | 25 | 25 | 45 | 3 |
| Initial cut high | 120 | 120 | 145 | 30 |
| Size low | 180 | 250 | 80 | 30 |
| Size high | 260 | 450 | 180 | 80 |

### SAMPL validation: measuring postural and locomotor kinematics in real-time

Next, to demonstrate how SAMPL facilitates efficient collection of high-quality kinematic data, we gathered a new dataset from larval zebrafish (7-9 days post-fertilization, dpf) that swam freely in the dark. A typical experimental repeat consisted of two sequential 24-hour sessions using 3 SAMPL boxes. Data were pooled across 27 repeats for subsequent analysis of kinematics. Each 24-hour behavior session yielded on average 1223±481 bouts per day for the standard chamber (6-8 fish) and 1251±518 bouts per day for the narrow chamber (1-3 fish). While not analyzed, running a single fish in the narrow chamber yielded 891±903 bouts over 24hrs. Based on the number of apparatus used, we estimate that a similar dataset (total n=121,979 bouts) could be collected in **two weeks** using a single SAMPL rack.

We first used our data to establish basic distributions of locomotion and posture. We used SAMPL’s processing algorithm to extract the following information in real-time: (1) pitch, defined as the angle between the long axis of the fish’s body and the horizon (Figure 2E); (2) x (azimuth), z (elevation) coordinates of the center of the pixels that correspond to the fish. After collection, we used SAMPL’s processing suite to extract basic postural kinematics during swimming. Zebrafish larvae swim in discrete periods of translation called “swim bouts” (Figure 2F) ^16, 20^. We defined swim bouts as periods where the instantaneous speed exceeds 5 mm/sec (Figure 2F, dashed line). The time of the peak speed was defined as t = 0 ms (Figure 2F, cyan lines). Swim bouts were aligned to peak speed for extraction of kinematic parameters; the period 250 ms before and 200 ms after peak speed was reserved for future analysis. We observed that zebrafish larvae swim predominantly at slower speeds with mean and standard deviation measured 12.90±4.91 mm/s, on par with previous reports ^16, 20–22^. Larvae showed a broad distribution of postures evaluated at peak speed (8.48°±15.23°) with a positive (nose-up) average, suggesting that SAMPL detected a variation of nose-up and nose-down swim bouts. SAMPL can thus rapidly acquire a rich dataset of spontaneous locomotor behavior and a wide range of “natural” postures.

### SAMPL validation: extracting key parameters of balance and vertical navigation in zebrafish

SAMPL includes data analysis and visualization code (Python source and sample datasets provided) optimized to extract key parameters of balance and locomotion from larval zebrafish.

We use our “two-week” dataset to demonstrate that SAMPL can resolve these four parameters:

Figure 3: Control of movement timing. ^16^

**Figure 3:**
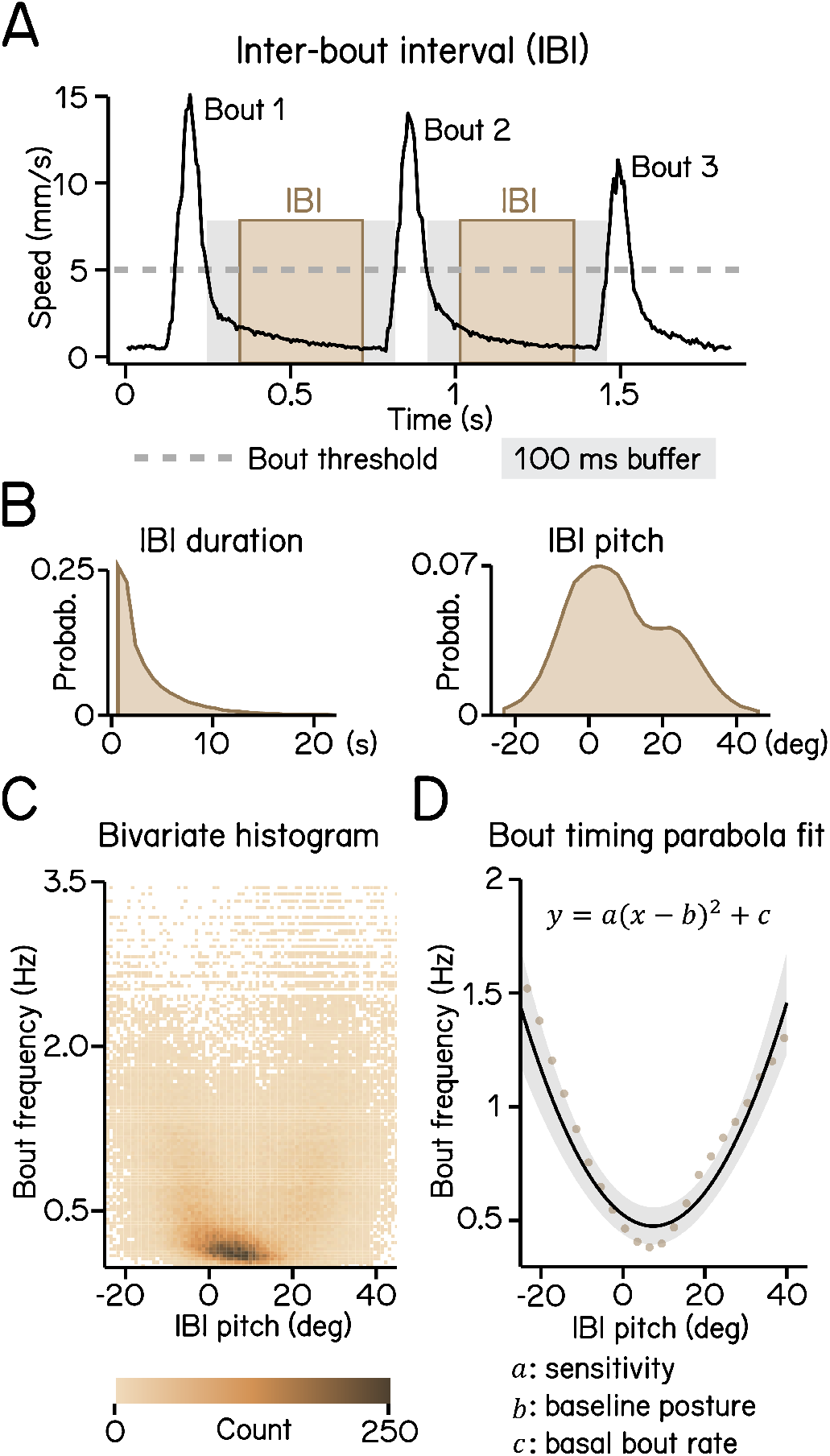
Modeling timing of swim bouts reveals larval sensitivity to pitch changes. (A) An inter-bout interval (IBI, brown area) is defined as the duration when swim speed is below the 5 mm/s homeostasis threshold (dashed line) between two consecutive bouts with a 100 ms buffer window (grey area) deducted from each end. (B) Distribution of IBI duration (left) and mean pitch angle during IBI (right). (C) Bivariate histogram of bout frequency and IBI pitch. Bout frequency is the reciprocal of IBI duration. (D) Bout frequency plotted as a function of IBI pitch and modeled with a parabola (black line, R^2^ = 0.14). Brown dots indicate binned average of IBI pitch and bout frequencies calculated by sorting IBI pitch into 3°-wide bins. For all panels, n = 109593 IBIs from 537 fish over 27 repeats.

Figure 4: Control of steering to climb/dive. ^17^

**Figure 4:**
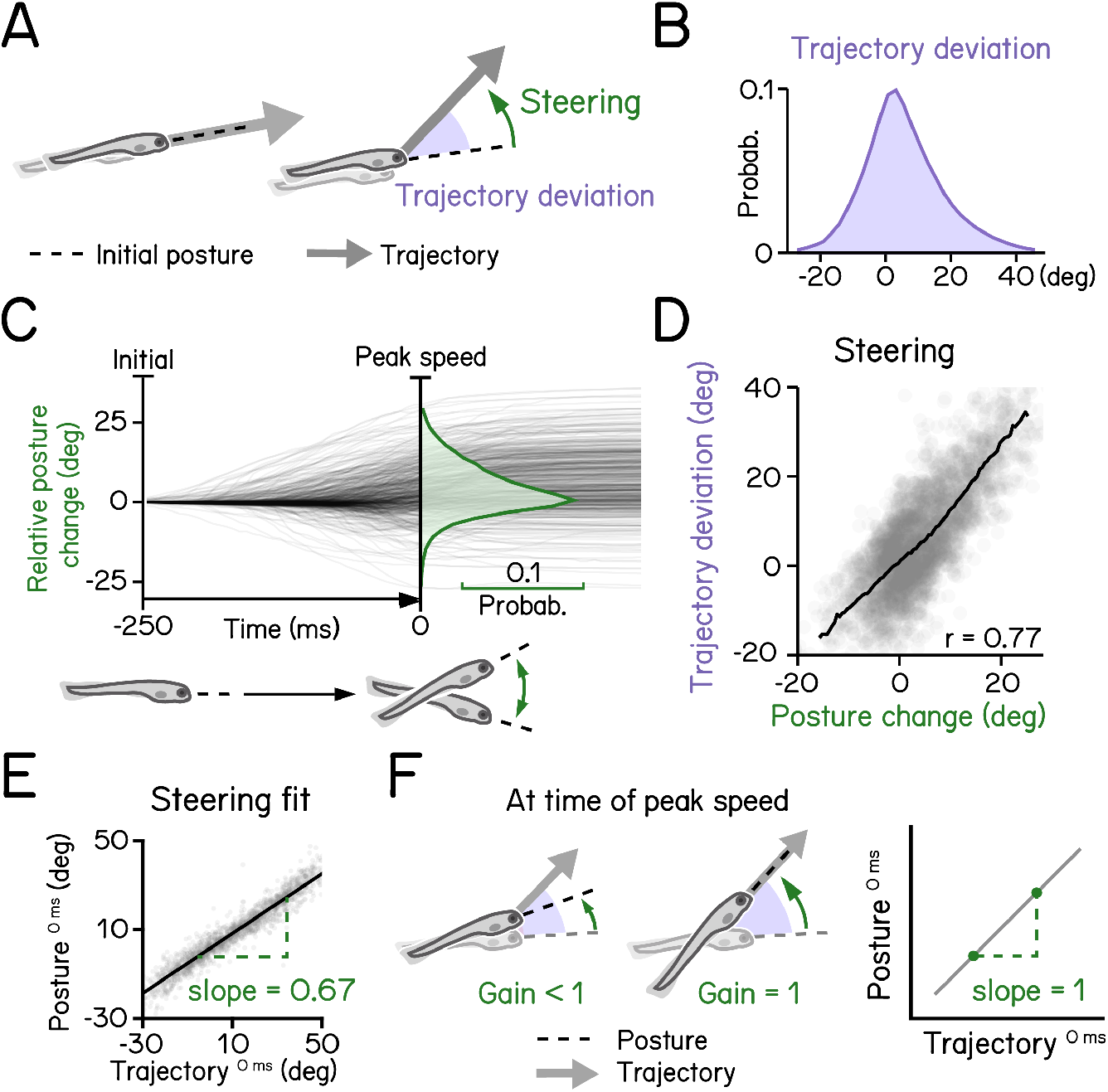
Larval vertical navigation is led by steering toward trajectory. (A) Schematic illustration of two climbing mechanics: (1) a larva may generate a thrust (arrow) toward the pointing direction (dashed line) at the initial of a bout (left); (2) a larva can steer (green arrow) toward an eccentric angle before the thrust (right). The offset between trust angle and the direction the larva point toward at bout initial is termed trajectory deviation (purple). (B) Distribution of trajectory deviation. (C) Changes of pitch angles relative to initial pitch plotted as a function of time (dark lines) overlaid with distribution of pitch change at time of peak speed (green). (D) Trajectory deviation (purple) plotted as a function of posture changes from bout initial to time of the peak speed (green). Black line indicates binned average values. Positive correlation between trajectory deviation and posture change demonstrates that larvae steer toward the trajectory of the bout. (E) To measure the gain of steering compared to trajectory deviation, pitch angels at time of the peak speed are plotted as a function of trajectory. Steering gain is determined as the slope of the fitted line (Pearson’s r = 0.96). (F) Schematic illustrations demonstrating how steering gain associates steering (green arrows) with trajectory deviation (purple). For all panels, n = 121979 bouts from 537 fish over 27 repeats.

Figure 5: Coordination between trunk and fin. ^18^

**Figure 5:**
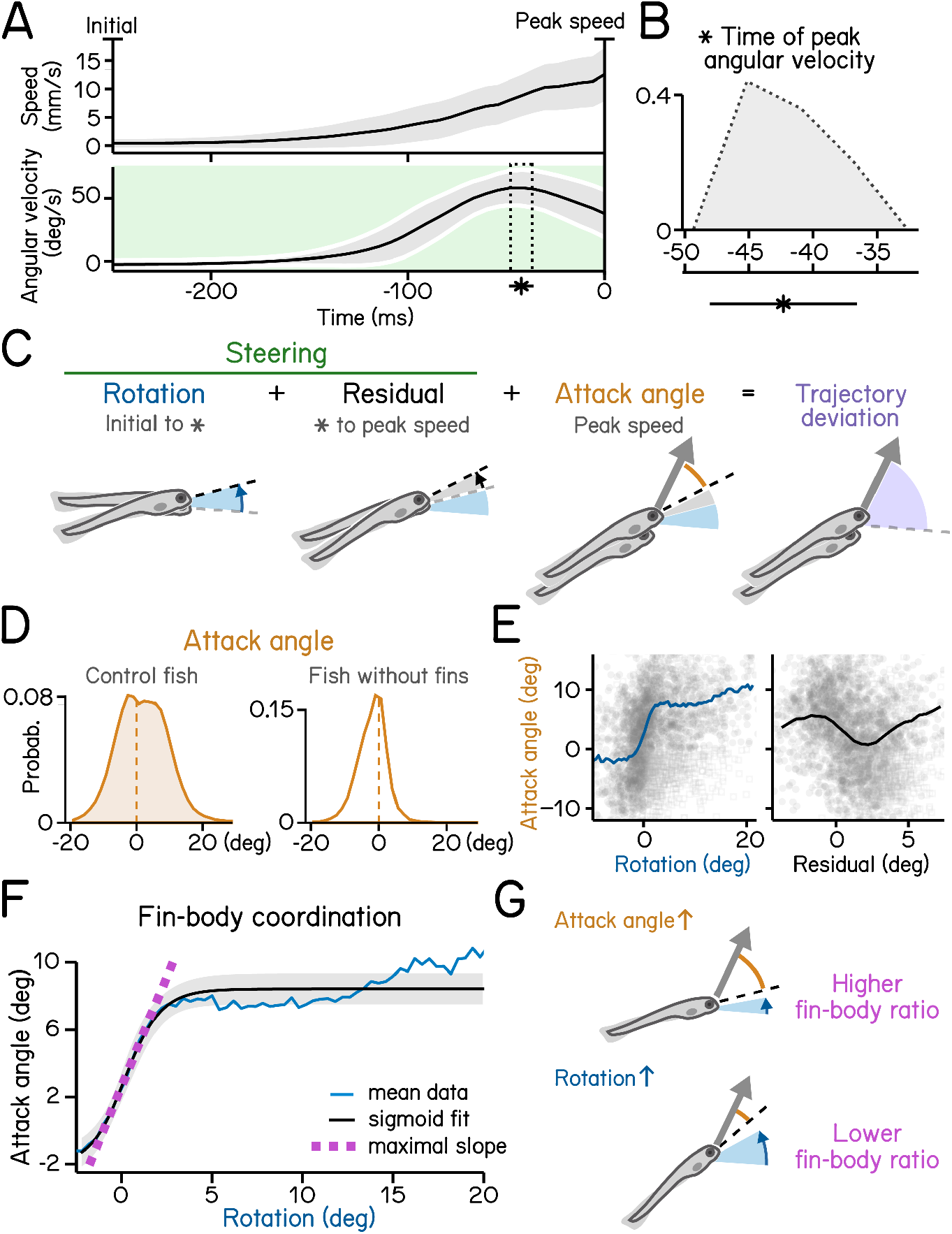
Steering requires coordination of fin and body. (A) Swim speed (top) and angular velocity (bottom) plotted as a function of time. Angular velocity peaks (asterisk and dotted area, mean SD) during steering phase (green) before time of the peak speed. Angular velocity is adjusted by flipping signs of bouts with nose-down rotations during steering (mean SD across experimental repeats). Shaded region in the upper panel indicates mean SD across all quantified swim bouts. (B) Histogram of time of peak angular velocity, binned by frame, across experimental repeats with mean SD plotted below. (C) Illustration of components that contribute to trajectory deviation. Larvae rotate their bodies starting from bout initial (blue) and reach peak angular velocity (asterisk) before peak speed. Any rotation generated during decrease of angular velocity is considered residual (grey). At time of peak speed, there is an offset between the pitch angle (dashed line) and bout trajectory (arrow) which is termed attack angle (orange). Body rotations, residual, and attack angle add up to trajectory deviation. (D) Distribution of attack angles in control fish (left) and fish after fin amputation (right). Dashed lines indicate 0 attack angle. (E) Attack angles plotted as a function of body rotations (left, blue) or residual rotations (right). Rotations and residuals are sorted into 0.5°-wide bins for calculation of binned average attack angles. Swim bouts with negative attack angles while having steering rotations greater the 50th percentile (hollow squares) were excluded for binned-average calculation. (F) Attack angles plotted as a function of body rotations (blue line) and fitted with a logistic model (black line, R^2^ = 0.31). Fin-body ratio is determined by the slope of the maximal slope of the fitted sigmoid (magenta). Rotations are sorted into 0.8°-wide bins for calculation of binned average rotations and attack angles (blue line). Swim bouts with negative attack angles while having steering rotations greater the 50th percentile were excluded for sigmoid modeling. (G) Schematic illustration of how fin-body ratio reflect climbing mechanics. For all panels, n = 121979 bouts from 537 fish over 27 repeats.

Figure 6: Control of posture stabilizing rotations. ^17^

**Figure 6:**
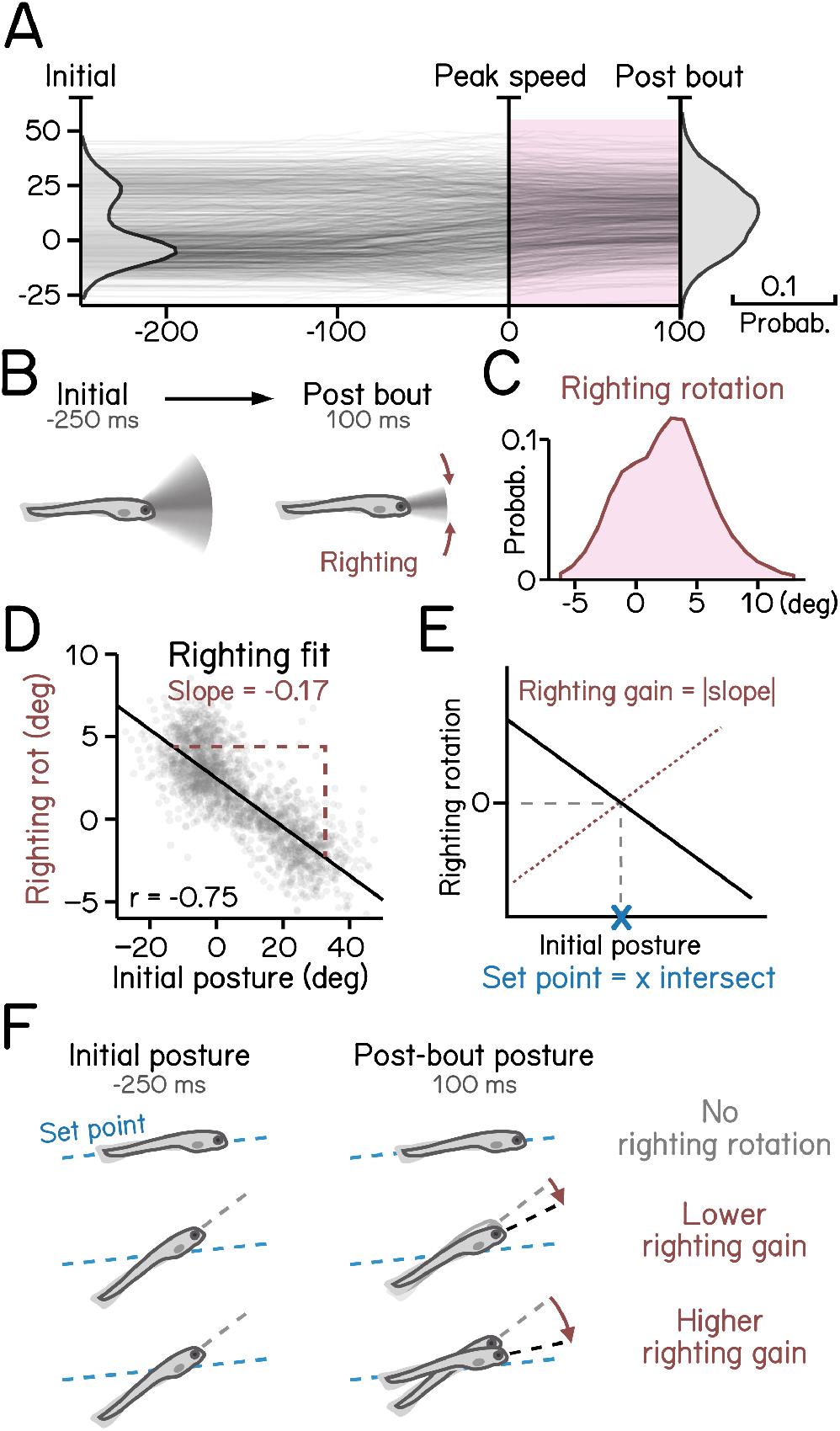
Righting rotation restores posture after peak speed. (A) Pitch angles plotted as a function of time (dark lines) overlaid with distribution of pitch angles before (left) and after bouts (right). Red area indicates duration after peak speed when pitch distribution narrowed. (B) Illustration of righting behavior. Larvae rotate (red arrows) toward more neutral posture after peak speed. (C) Distribution of rotation during righting (red in **A**). (D) Righting rotation plotted as a function of initial pitch angles. (E) Righting gain is determined by the absolute value of the slope (red dotted line) of best fitted line (black line). The x intersect of the fitted line determines the set point (blue cross) indicating posture at which results in no righting rotation. (F) Schematic illustration of righting rotation (red arrows), righting gain, and set point (blue dashed line). For all panels, n = 121979 bouts from 537 fish over 27 repeats.

We conclude that SAMPL’s resolution and throughput allows rapid and deep insight into each parameter, detailed below. Data analysis using the provided scripts on the provided dataset runs in 30 minutes on a typical analysis computer (M1 processor, 16GB RAM). Full details of analysis/visualization is provided in Appendix 4, and a step-by-step guide to set up the relevant environment and to run experiments provided in Appendix 5.

Proper balance requires active stabilization. Zebrafish larvae are front-heavy and therefore subject to destabilizing torques in the pitch (nose-up/nose-down) axis. Swim bouts counteract the resultant forces, stabilizing the fish. Zebrafish larvae learn to initiate swim bouts when unstable ^16^. We first defined movement rate as the reciprocal of the inter-bout interval (Figures 3A and 3B). More extreme postures were associated with higher movement rate (Figure 3C), with a parabolic relationship (Figure 3D, R^2^ = 0.14). We expect that the majority of the residual variance reflects a previously-reported dependence of movement timing on angular velocity ^16^. The three coefficients of the parabola represent the baseline posture, the basal rate of movement, and – key to our analysis – the degree to which postural eccentricity relates to movement rate, or “sensitivity,” (Figure 3D). SAMPL therefore permits efficient quantification of a crucial posture-stabilizing behavior: the relationship between perceived instability and corrective behavior.

Like most animals, larval zebrafish go where their head points. To adjust their vertical trajectory (i.e. to climb or dive) larvae must rotate their bodies away from their initial posture, pointing in the direction they will travel (Figures 4A and 4B) ^17, 23^. Previous work ^17^ established that steering rotation in larvae swimming spontaneously occurs mostly before they reach the peak speed (Figure 4C). A larva’s steering ability reflects the relationship between the change in posture before the peak speed and the resultant deviation in trajectory (Figure 4D). We parameterized steering as the slope (gain) of the best-fit line between posture and trajectory evaluated at the time of peak speed (Figure 4E). A gain of 1 indicates that the observed trajectory could be explained entirely by the posture at the time of peak speed (Figure 4F). SAMPL revealed that 7 dpf larvae exhibit an average steering gain at 0.67, suggesting an offset between posture and trajectory at the time of peak speed (Figure 4E, R^2^ = 0.92). SAMPL allows us to infer how effectively larvae steer using axial (trunk) musculature to navigate the water column.

To climb (Figures 5A and 5B) fish generate lift with their pectoral fins, assisting steering rotations and subsequent axial undulation ^24, 25^. Larval zebrafish learn to climb efficiently by coordinating their trunk and fins ^18^. We defined the attack angle, or the additional lift associated with each climb, as the difference between the steering-related changes and the resulting trajectory (Figure 5C). We evaluated attack angle after pectoral fin loss, revealing a clear contribution to climbs (Figure 5D). Next, we demonstrate a positive correlation (with rectification and asymptote) between steering-related rotations and fin-based attack angle (Figure 5E, left). Notably, after peak angular velocity, rotations are poorly correlated with attack angles (r = −0.17) (Figure 5E, right). These residuals reflect the initial angular deceleration as fish reach their peak speed (Figure 5A). We parameterize the relationship between the initial rotation and the attack angle using logistic regression (Figure 5F, R^2^ = 0.31). The regression reveals the maximal slope of the sigmoid relating steering and lift (Figure 5G). We named this slope “fin-body ratio” as it describes how larvae distribute labor between axial and appendicular muscles, i.e. between trunk (steering) and fins (lift), as shown in previous work ^18^. SAMPL thus permits efficient inference of coordinated behavior.

Larvae must actively maintain their preferred posture in the pitch axis. To do so, they rotate partially towards their preferred orientation as they decelerate (Figures 6A to 6C). The magnitude of these rotations scales with the eccentricity of their posture before a swim bout ^17^. We estimated the slope (−0.17) of the line that related initial posture and the amount the fish rotated back toward the horizontal (Figure 6D), R^2^ = 0.56. As the behavior is corrective, the relationship is negative; we therefore define the gain of righting as the inverse of the slope (Figure 6E). We further define the “set point” as the point where an initial posture would be expected to produce a righting rotation of zero (Figures 6E and 6F). SAMPL facilitates quantification of corrective reflex abilities (gain) and associated internal variables (set point).

Taken together, our estimates of key posture and locomotor parameters establish that SAMPL can rapidly generate datasets that permit rich insight into the mechanisms of balance and vertical navigation.

### SAMPL can resolve slight variations in posture control strategies across genetic backgrounds

To be useful SAMPL must resolve small but systematic differences in key measures of posture and vertical locomotion. Even among isogenic animals reared in controlled environments, genetic differences contribute to behavioral variability ^26–33^. The “two-week” dataset analyzed in Figures 3 to 6 included data from three different genetic backgrounds. Larvae for experiments were generated by crossing the same clutch of wild-type adults (mixed background) to zebrafish of three different strains: AB (n = 62457 bouts, N = 225 fish over 10 experimental repeats); SAT (n = 27990 bouts, N = 117 fish over 7 experimental repeats); and the lab wild type (n = 31532 bouts, N = 195 fish over 10 experimental repeats), which resembles real-world approaches where a key transgenic line is often crossed to different backgrounds for experiments. To capture the full variance in the dataset, we took a conservative approach by calculating kinematic parameters for individual experimental repeats (n = 4518±1658 bouts). We assayed SAMPL’s sensitivity by asking (1) if there were detectable differences in the four parameters defined in Figures 3 to 6 and (2) if these differences were systematic.

Qualitatively, larval zebrafish of the same age swim similarly; as expected, the magnitude of change across strains we observed in Figure 7 is quite small. Nonetheless SAMPL could resolve systematic variations in locomotion behavior and balance abilities among larvae of different strains (Figure 7). AB larvae exhibited the best posture stability, demonstrated by the lowest standard deviation of IBI pitch compared to the other two strains (Figure 7C). Correspondingly, AB larvae had the highest bout frequency (Figure 7B), sensitivity to posture changes (Figure 7E), and righting gain (Figure 7K), all of which contributes positively to their higher posture stability. These results demonstrate that SAMPL is capable of detecting inter-strain variations in locomotion and balance behavior.

**Figure 7:**
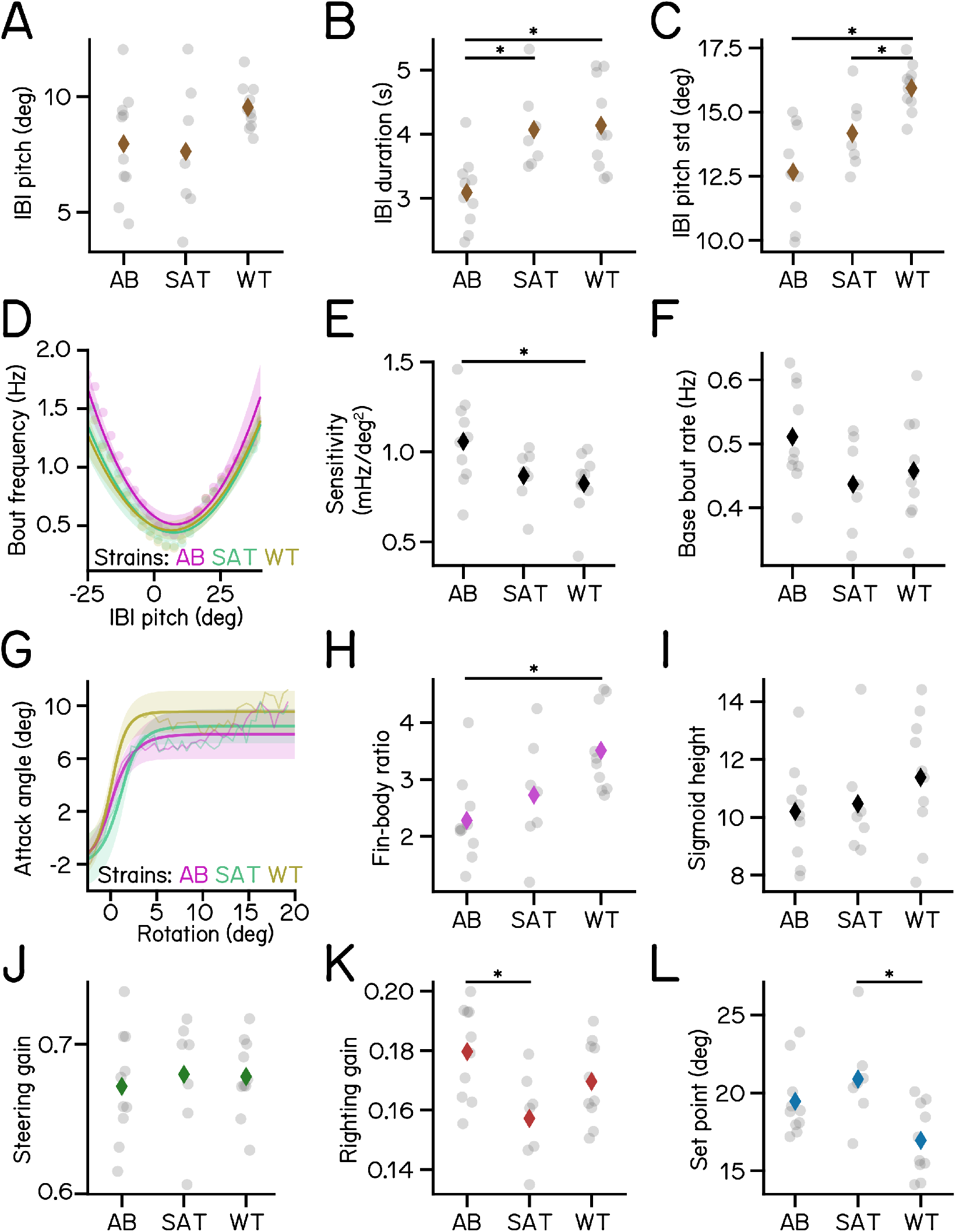
Variations of kinematic parameters among three different zebrafish strains. (A) Average pitch angles during IBI. (B) IBI duration (AB vs SAT p-adj = 0.0128; AB vs WT p-adj = 0.0034). (C) Standard deviation of IBI pitch (AB vs WT p-adj = 0.0001; SAT vs WT p-adj = 0.0479). (D) Bout frequency plotted as a function of IBI pitch modeled with parabolas. (E) Sensitivity to pitch changes (AB vs WT p-adj = 0.0319). (F) Baseline bout rate. (G) Attack angles plotted as a function of body rotations modeled with sigmoids. (H) Fin-body ratio (AB vs WT p-adj = 0.0066). (I) Height of the sigmoid in **G**. (J) Steering gain of different strains. (K) Righting gain of different strains (AT vs SAT p-adj = 0.0133). (L) Set point (SAT vs WT p-adj = 0.0094). For each strain of AB/SAT/WT, N = 10/7/10 repeats, n = 62457/27990/31532 bouts and 55683/25964/27946 IBIs from 225/117/195 fish.

In contrast, larvae of different ages adopt different strategies to stabilize posture and navigate in depth ^16–18^ To contextualize the magnitude of strain-related differences we gathered a longitudinal dataset by measuring behavior from the same siblings of the AB genotype at three timepoints: 4-6, 7-9, and 14-16 dpf (Table 3). We observed that the standard deviation of IBI pitch for 4 and 14 dpf larvae was 38.1% higher and 11.3% lower, respectively, than the average result of 7 dpf larvae (Table 3). Across strains at 7 dpf, the variation was much smaller: from 11.8% higher to 11.2% lower. Similarly, relative to 7 dpf larvae, sensitivity of 4 dpf larvae was considerably lower (−42.5%), and increased to 23.6% higher by 14 dpf (Table 3); variations among 7 dpf strains were up to 10.0% lower and 15.4% higher.

**Table 3:** Measured parameters of posture and locomotion across development

| Parameter | Unit | 4 dpf | 7 dpf | 14 dpf | Format | Definition |
| --- | --- | --- | --- | --- | --- | --- |
| Peak speed | mm/s | 10.42<br>(3.85) | 13.02<br>(4.99) | 11.41<br>(4.20) | Mean of bouts (SD) | Peak speed of swim bouts |
| Initial pitch | deg | 5.21<br>(31.49) | 0.77<br>(21.81) | 0.54<br>(18.64) | Median of bouts (IQR) | Pitch angle at 250 ms before the peak speed |
| Pitch at peak speed | deg | 9.74<br>(29.16) | 6.84<br>(20.35) | 4.36<br>(19.73) | Median of bouts (IQR) | Pitch angle at time of the peak speed |
| Post-bout pitch | deg | 10.57<br>(23.86) | 10.21<br>(16.70) | 6.88<br>(16.05) | Median of bouts (IQR) | Pitch angle at 100 ms after the peak speed |
| End pitch | deg | 10.85<br>(23.59) | 10.78<br>(16.79) | 7.56<br>(15.55) | Median of bouts (IQR) | Pitch angle at 200 ms after the peak speed |
| Bout trajectory | deg | 12.29<br>(27.52) | 8.92<br>(20.19) | 7.85<br>(22.89) | Mean of bouts (SD) | Peak trajectory, tangential angle of the trajectory at the time of the peak speed |
| Bout displacement | mm | 1.12<br>(0.63) | 1.36<br>(0.64) | 1.35<br>(0.70) | Mean of bouts (SD) | Average displacement of fish during a bout when speed is greater than 5 mm/s |
| Inter-bout interval | s | 1.78<br>(2.61) | 1.89<br>(2.75) | 2.13<br>(2.80) | Median of bouts (IQR) | IBI, duration between two adjacent swim bouts |
| Bout frequency | Hz | 0.56<br>(0.70) | 0.53<br>(0.69) | 0.47<br>(0.59) | Median of bouts (IQR) | Frequency of swim bouts determined by the reciprocal of inter-bout interval |
| IBI pitch | deg | 8.75<br>(17.73) | 8.06<br>(13.07) | 6.08<br>(11.24) | Mean of bouts (SD) | Mean pitch angle during inter-bout interval |
| IBI pitch standard deviation | deg | 17.48<br>(1.60) | 12.66<br>(1.80) | 11.23<br>(1.28) | Mean of repeats (SD) | Standard deviation of IBI pitch, a measurement of stability |
| Sensitivity | mHz/deg<sup>2</sup> | 0.61<br>(0.18) | 1.06<br>(0.23) | 1.31<br>(0.34) | Mean of repeats (SD) | Sensitivity to pitch changes. Determined by the coefficient of the quadratic term of the parabola model for bout timing |
| Baseline bout rate | Hz | 0.51<br>(0.06) | 0.51<br>(0.08) | 0.47<br>(0.11) | Mean of repeats (SD) | Y intersect of the parabola model for bout timing |
| Trajectory deviation | deg | 5.46<br>(14.53) | 4.13<br>(11.57) | 4.35<br>(16.39) | Mean of bouts (SD) | Deviation of bout trajectory from initial pitch |
| Steering rotation | deg | 2.30<br>(7.51) | 3.00<br>(7.33) | 1.94<br>(6.31) | Mean of bouts (SD) | Change of pitch angle from initial (250 ms before) to the time of the peak speed |
| Steering gain | - | 0.64<br>(0.04) | 0.67<br>(0.04) | 0.51<br>(0.05) | Mean of repeats (SD) | Slope of best fitted line of posture vs trajectory at the time of the peak speed |
| Steering-related rotation | deg | 1.72<br>(6.15) | 1.74<br>(5.95) | 0.99<br>(5.42) | Mean of bouts (SD) | Change of pitch angle from initial to the time of max angular velocity |
| Attack angle | deg | 4.10<br>(16.16) | 0.77<br>(9.68) | 0.91<br>(5.25) | Median of bouts (IQR) | Deviation of bout trajectory from pitch at time of the peak speed |
| Peak angular velocity time | ms | 50.60<br>(7.62) | 39.16<br>(4.96) | 50.00<br>(5.12) | Mean of repeats (SD) | Time of peak angular velocity in ms before time of the peak speed |
| Fin-body ratio | - | 3.41<br>(0.86) | 2.27<br>(0.76) | 3.55<br>(1.98) | Mean of repeats (SD) | Maximal slope of best fitted sigmoid of attack angle vs early rotation |
| Sigmoid height | deg | 16.47<br>(2.31) | 10.28<br>(1.78) | 25.15<br>(4.68) | Mean of repeats (SD) | Height of best fitted sigmoid of attack angle vs early rotation |
| Righting rotation | deg | 0.92<br>(3.49) | 2.64<br>(3.55) | 1.90<br>(3.01) | Mean of bouts (SD) | Change of pitch angle from time of the peak speed to post bout (100 ms after peak speed) |
| Righting gain | - | 0.15<br>(0.02) | 0.18<br>(0.02) | 0.18<br>(0.02) | Mean of repeats (SD) | Numeric inversion of the slope of best fitted line of righting rotation vs initial pitch |
| Set point | deg | 13.00<br>(2.10) | 19.47<br>(2.28) | 13.60<br>(1.78) | Mean of repeats (SD) | X intersect of best fitted line of righting rotation vs initial pitch |

Our analysis of new data supports three key conclusions. First, SAMPL can uncover small, systematic differences in the way fish swim and stabilize posture. Second, SAMPL can make longitudinal measures of the same complement of animals as they develop. Third, relative to development, the effect of genetic background is small. We conclude that SAMPL’s capacity to resolve small differences supports its usefulness as a tool screen for modifiers of postural control and vertical locomotor strategies.

### Estimating SAMPL’s resolution

Our dataset establishes SAMPL’s ability to resolve small kinematic differences between cohorts. How does SAMPL’s power change as a function of the size of the dataset? We used resampling statistics to estimate SAMPL’s resolution as a function of the number of the bouts (Methods). To ensure our most conservative estimate, we resampled data combined across AB, SAT and WT genotypes at 7dpf.

As expected, the width of the confidence interval for any estimated parameter decreased with the number of bouts (Figure 8A). The most challenging parameter to estimate is coordination between fin and trunk (fin-body ratio) The steepness with which the confidence interval width decreases follows the number of regression coefficients necessary for each measure: fin-body ratio (4 parameters); bout timing (3 parameters); and steering or righting (2 parameters). We therefore propose that these particular measures can serve as a general guide for the challenge of estimating parameters within a SAMPL dataset.

**Figure 8:**
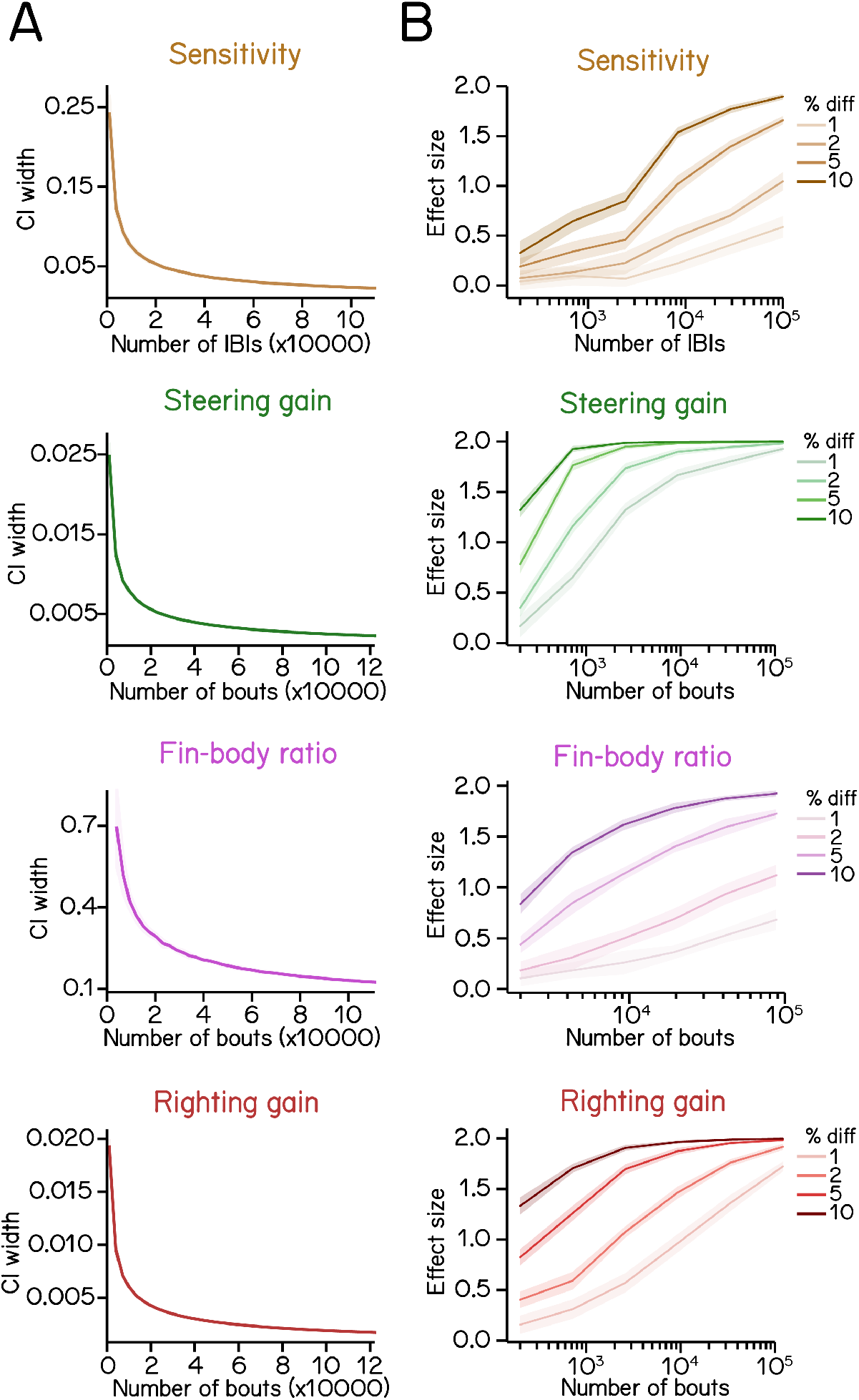
Statistics of regression analysis for swim kinematics. (A) Confidence interval (CI) width of kinematic parameters plotted as a function of sample size at 0.95 significance level (mean ± SD as ribbon). Errors were estimated by resampling with replacement from the complete dataset. (B) Effect size plotted as a function of sample size at various percentage differences. Refer to Methods for details of computation.

A fundamental challenge for all screens is determining the sample size required to correctly reject the null hypothesis ^34^. We address this question by asking how much data one would need to gather in order to detect meaningful effects. We simulated difference of particular magnitudes by imposing an offset on each parameter (sensitivity, steering gain, fin-body ratio, and righting gain) while preserving the original variance (Methods). Offsets were expressed as a fractional difference, and resampling was used to estimate the effect size one would see as a function of the number of bouts/IBIs when comparing kinematic parameters between the original dataset and the dataset with an imposed effect (Methods).

Broadly, we find that for all kinematic parameters, the smaller the percent change, the larger the required sample size (Figure 8B). Steering and righting gains require the fewest bouts to detect a 1-2% change with an effect size > 0.5 (Figure 8B, green and red). However, sensitivity and fin-body ratio require relatively larger datasets to confidently discriminate small changes (Figure 8B, brown and magenta). We conclude that the full “two-week” dataset we generated using SAMPL (n = 121,979 bouts) is sufficient to reveal any biologically-relevant differences between two conditions.

In summary, these simulations demonstrate that a single SAMPL rack divided into two conditions (6 apparatus / each) could, in two standard 48-hour runs, generate sufficient data to resolve meaningful differences in postural and locomotor kinematics between two conditions. We provide detailed instructions in Appendix 5 addressing experimental design strategies to maximize SAMPL’s resolution.

## DISCUSSION

We present SAMPL, a scalable solution to measure posture and locomotion in small, freely-moving animals. We start with a brief overview of the hardware and software, with comprehensive guides to every aspect of SAMPL’s hardware and software included in the Appendices. Next we illustrate SAMPL’s flexibility with raw video & real-time measurements from three common model organisms: *Drosophila melanogaster* (fly), *Caenorhabtitis elegans* (worms), and *Danio rerio* (zebrafish). To illustrate the depth of insight accessible using SAMPL we explored a new dataset – consisting of two weeks worth of data – that illuminates four key parameters of zebrafish navigation in depth: bout timing, steering, fin-body coordination, and righting. We made two discoveries using SAMPL’s analysis suite: (1) systematic changes to zebrafish posture and locomotion across genetic backgrounds and (2) that these changes were small relative to variation across developmental time. Finally, we use our new dataset to define SAMPL’s resolution: how much data an experimenter would need to collect to detect meaningful effects. Taken together, SAMPL provides a screen-friendly solution to investigate vertical locomotion and/or other behaviors using common small model organisms, and a turn-key solution to study balance in larval zebrafish. More broadly, our approach serves as a template for laboratories looking to develop or scale their own hardware/software. Below we detail SAMPL’s innovations and limitations, and make a case for screens to address unmet clinical needs for balance disorders.

### SAMPL’s innovations

One of SAMPL’s key innovations is to measure vertical behavior, where the effects of gravity play a role. The overwhelming majority of studies monitor animal behavior from above, where animals are constrained to a horizontal plane. For most animals – especially those that swim or fly – vertical navigation and its neuronal representation ^35, 36^ is vital. Further, maintaining posture in the face of gravity is a universal challenge ^37–39^, particularly as animals develop ^16, 40^. SAMPL can illuminate animal trajectories during exploration of depth.

SAMPL reduces the dimensionality of behavior along a number of axes in real-time. First, by focusing on a homogeneous part of the behavioral arena, SAMPL bypasses a number of imaging challenges and difficulties involved in interpreting behavior along arena walls ^41^. Second, by rejecting frames with multiple animals in view at the same time SAMPL incorporates animal-to-animal variability ^4^ within each estimated parameter without having to keep track of individuals; the narrow chamber (Figure 1E) is ideal for single-animal experiments if such variability is of interest. Third, while large enough to permit unconstrained behavior, the anisotropic dimensions of SAMPL’s behavioral arenas (Figure 1E) facilitate measurements in the vertical axis. SAMPL’s design choices thus facilitate rapid extraction of behavioral parameters relevant for posture and locomotion.

SAMPL was designed to scale efficiently. Data is gathered by a compiled executable, allowing SAMPL to run three apparatus off a single computer, reducing costs and space. A SAMPL rack consists of 12 apparatus running off four computers with a footprint of 24”x36”x81.5” (LxWxH). The key components such as the camera are readily available from multiple suppliers. Taken together, SAMPL can be used immediately to screen and/or to provide videographic data from freely moving animals at scale.

Our new dataset, gathered in two weeks, illustrates the power of SAMPL’s analysis/visualization workflow for studies of larval zebrafish balance. While SAMPL can and does save video, by sign it extracts only three parameters in time: the (x,z) coordinates of the animal and the angle between the body and the horizon. As we demonstrate here, this small set of parameters defines behaviors larval zebrafish use to swim and balance in depth: bout timing (Figure 3), steering (Figure 4), fin-body coordination (Figure 5), and righting (Figure 6). While each parameter has been previously defined ^16–18^, the new data we present here illustrates differences across genetic backgrounds and development and allows granular estimation of statistical sensitivity. Taken together, SAMPL’s focus facilitates exploration of unconstrained vertical behavior.

### Comparisons with other approaches

Here, we discuss SAMPL’s advantages by comparing it with other available tools for measuring *Drosophila*, *C. elegans*, and zebrafish behavior.

### SAMPL for measuring *Drosophila* behavior

SAMPL offers advantages over previous methods for measuring negative gravitaxis, an innate behavior of *Drosophila melanogaster* ^42^. The most widespread method, called the bang test, consists of banging flies down inside a vertical tube and then counting the number of flies that walk an arbitrary vertical distance in an arbitrary amount of time ^42–45^. This method startles the flies, which may confound the behavior, and the flies are limited in directional choice. Using SAMPL, a measurement of fly vertical position and orientation is instantaneously acquired without needing to startle the flies. Another *Drosophila* gravitaxis assay is the geotaxis maze ^46^, that allows the flies to make a series of up-or-down choices as they move across the maze towards a light. While the flies are not startled in this assay, they are still constrained to moving only up or down. SAMPLs high resolution camera permits continuous monitoring of free vertical walking behavior, as well as high-resolution monitoring of head, wing, leg, and antenna positions. While SAMPL has been designed to monitor behavior in the vertical plane, the hardware and software strategies we have developed for high throughput recording could be similarly adapted to increase the throughput of measuring other *Drosophila* behaviors such as grooming ^47^, sleep ^48^, courtship ^49^, and aggression ^50^. Because SAMPL has both high resolution recording and the ability to scale, screening through microbehaviors like head tilting or limb positioning is possible. Notably, an earlier version of SAMPL’s detection algorithm was successfully used for data acquisition in a fly olfactory behavior assay ^51, 52^ with minimal changes. Taken together, SAMPL’s resolution, throughput, and adaptability complement and extend current approaches to measure *Drosophila* behavior, particularly in the vertical axis.

### SAMPL for measuring *C. elegans* behavior

The simple nervous system of *C. elegans* is a powerful model to study neural circuits that control posture and movement. *C. elegans* possess a rich and tractable repertoire of motor control ^53^. For example a pattern generator creates sinusoidal waves of muscle contraction that propel *C. elegans* on a solid substrate, and these sinusoidal movements are sculpted by proprioceptive feedback ^54^. Proprioceptive feedback also controls transitions between sinusoidal crawling and non-sinusoidal bending that can propel animals in a liquid environment ^55–57^.

Other sensory stimuli elicit coordinated motor responses that are critical for navigation. De- creasing concentrations of attractive odorants and gustants trigger reversals followed by a pirou- ette or omega bend, which results in a large-angle turn that reorients animals ^58, 59^. A distinct navigation behavior involves precise steering of an animal as it follows an isotherm in a tem- perature gradient ^60, 61^ or tracks a preferred concentration of gustant ^62^. The resolution and scalability of SAMPL offers the opportunity to determine the cellular, molecular, and genetic underpinnings of these diverse motor control mechanisms.

*C. elegans* behavior becomes complex in enriched 3D environments, with animals using strate- gies for exploration and dispersal not seen under standard laboratory conditions ^63^. Behavior trackers that have been used to study *C. elegans* kinematics are generally restricted to analysis of behaviors on a surface. By contrast, SAMPL measures behavior in a volume and is well-suited to the study of newly discovered behaviors that are only expressed in environments that vary across depth. One such example is gravitaxis, where *C. elegans* display both positive ^64^ and negative gravitaxis ^65^, underscoring the need for additional pipelines to test behavior ^66^. The new data we present here establishes that SAMPL offers a powerful complement to existing pipelines for *C. elegans* assays of behavior in the vertical dimension.

### SAMPL for measuring zebrafish behavior

SAMPL joins a decades-long tradition of apparatus that has, collectively, established the larval zebrafish as a key vertebrate model to understand the neural control of posture and locomotion ^13–15^. Broadly, these devices sit on a continuum that represents a trade-off between imaging resolution and throughput. At one end, exquisite measures of tail or eye kinematics are available when imaging single animals that are partially restrained ^67^, or contained in a small field of view ^68^. Such devices are particularly useful when combined with imaging or perturbations of neuronal activity, but at the cost of throughput. At the other end are devices that measure activity when single animals are constrained to small arenas, such as the ∼8 mm^2^ wells in a 96-well plate ^6,69–71^. These devices lend themselves well to screens, and offer commercial options, but the range of behaviors is compressed ^72^. Like other attempts to preserve high-resolution kinematic information while accommodating natural unconstrained behavior ^22, 73–78^, SAMPL sits between these two extremes, joining other open-source software packages such as Stytra ^79^ and Zebrazoom ^80^. We see SAMPL as a complementary tool. SAMPL’s emphasis on vertical behavior and its scalability position it to leverage the advantages of the zebrafish model for screens – either as a primary resource, or to follow-up on promising “hits” identified with higher-throughput approaches ^6^.

### Screening

Balance disorders present a profound and largely unmet clinical challenge ^19^. Because the neuronal architecture for balance is highly conserved and the fundamental physics (i.e. gravity is destabilizing) is universal, animal models represent a promising avenue for discovery. Due to their size, low cost, molecular accessibility, high fecundity, and conserved biology small animals – both vertebrates and invertebrates ^81^ – have long been used in successful screens of both candidate genes ^82^, peptides ^83^ and therapeutics ^84, 85^. Zebrafish are an excellent exemplar, particularly in the space of neurological disorders ^3^, with well-established approaches for candidate gene screens ^2, 5^, peptides ^86^, small molecules ^87–91^, and disease models ^92^. Using SAMPL with zebrafish, our dataset establishes a foundation to screen for balance modifiers in health & disease.

One particular arena where zebrafish screens for balance/posture could have a profound impact is in addressing the unmet therapeutic need that exists for a neurodegenerative tauopathy: progressive supranuclear palsy (PSP). PSP is initially characterized by balance impairments, falls, vertical gaze palsy, and rigidity ^93, 94^. Falls are central to early ^95^ PSP presentation and diagnosis ^96, 97^ and lead to fractures and hospitalization ^96, 98^. Currently, no treatments improve balance. Studies of posture ^99–103^, graviception ^104^, reflexes ^105–108^, electromyography ^109, 110^, and neural balance circuits in PSP ^103, 111–115^ are often underpowered, inconsistent, and have yet to identify the specific mechanism or substrate causing falls. Like most genes and subcortical structures ^116–123^ the genetic and anatomical substrates of PSP are conserved between humans and zebrafish ^124–127^. Here, using SAMPL, we define behavioral endpoints that reflect how pathological zebrafish might “fall.” By establishing SAMPL’s resolution, our data lay the foundation for impactful discovery in the space of a neurodegenerative disorder with balance pathology.

### Future prospects

SAMPL uses low-cost videographic and computing hardware to make novel behavioral measurements. By optimizing scalability, resolution, and extensibility, SAMPL allows experimenters to rapidly measure unconstrained behavior as animals navigate in depth. We have used SAMPL with a model vertebrate, zebrafish, to gain insight into posture and vertical locomotion, and to lay the groundwork for future screens. A wide variety of neurological disorders present with balance and locomotor symptoms. SAMPL offers a way to both understand the fundamental biology of balance, as well as means to evaluate candidate therapeutics to address this unmet need. More broadly, SAMPL stands as an exemplar and resource for laboratories looking to develop, adapt, or scale videographic apparatus to measure behavior in small animals.

### Limitations of the study

Any apparatus necessarily reflects a set of trade-offs. Consequentially, each of SAMPL’s innovations can reasonably be recast as a limitation depending on experimental priorities. For example, SAMPL’s focus on a subset of space and parameters is ill-suited to reconstruct a catalog of behaviors from videographic measurements i.e. a computational ethogram ^11, 20^. Similarly, SAMPL assumes that the animal’s trajectory reflects coordinated use of its effectors (limbs/trunk/wings). While SAMPL’s videos would be an excellent starting point for markerless pose estimation, detailing the links between effector kinematics and resultant changes to posture and trajectory may be better served by a multi-camera setup ^8, 9^. SAMPL’s processing is exclusive to one animal; other approaches are therefore necessary to resolve social interactions ^7, 128^. Finally, SAMPL’s analysis/visualization toolset incorporates priors for movement of zebrafish only – studies of other species would require a moderate investment of effort.

## Supporting information

Movie 1: SAMPL assembly Timelapse

Movie 2: SAMPL data from fly and worm

Movie 3: SAMPL data from fish

## ACKNOWLEDGMENTS

Research was supported by the National Institute on Deafness and Communication Disorders of the National Institutes of Health under award numbers R00DC012775 (DS), R56DC016316 (DS), R01DC017489 (DS), and F31DC019554 (KRH), and the National Institute of Neurological Disorders and Stroke under award numbers, T32NS086750 (KRH) and R61NS125280 (DS), by the Leon Levy Foundation (YZ, CEM), and the Rainwater Charitable Foundation (YZ) and by the Irma T. Hirschl/Monique Weill-Caulier Trust (DS). The authors thank David Ehrlich and members of the Schoppik and Nagel laboratories for their valuable feedback and discussions.

---

**Movie 1**

Movie 1. Stop motion instruction for box assembly.

**Movie 2**

Movie 2. Example of recorded epochs of a fly, a shrimp, and a worm. Scale bar: 2 mm.

**Movie 3**

Movie 3. Top: example of a recorded epoch of a freely-swimming zebrafish larva using the apparatus. Bottom: swim speed and pitch angles plotted as a function of time. Scale bar: 1 mm.

## STAR METHODS

### RESOURCE AVAILABILITY

#### Lead contact

Further information and requests for resources and reagents should be directed to and will be fulfilled by the lead contact, Dr. David Schoppik.

#### Materials availability

This study did not generate new unique reagents.

#### Data and code availability

SAMPL source code, SAMPL executables, raw behavior data, analyzed data used to make paper figures and README.md descriptions of each are all deposited with the Open Science foundation and are publicly available. DOI is listed in the key resources table. All original code has been deposited at the Open Science foundation and is publicly available. DOI is listed in the key resources table. This resource includes code to generate each figure / table in this manuscript. Any additional information required to reanalyze the data reported in this paper is available from the lead contact upon request.

### EXPERIMENTAL MODEL AND SUBJECT DETAILS

All procedures involving larval zebrafish (*Danio rerio*) were approved by the New York University Langone Health Institutional Animal Care & Use Committee (IACUC). Zebrafish larvae were raised at 28.5°C on a standard 14/10 h light/dark cycle at a density of 20-50 larvae in 25-40 ml of E3 medium before 5 days post-fertilization (dpf). Subsequently, larvae were maintained at densities under 20 larvae per 10 cm petri dish and were fed cultured rotifers (Reed Mariculture) daily. Larvae that had their behavior measured at 14 dpf were raised as stated above before being moved to 2 L tanks with 300 ml of cultured rotifers at 9 dpf. At 13 dpf, they were transferred back to petri dishes with E3 medium for adaptation.

Larvae with different strains were achieved by crossing Schoppik lab strain with a mixed AB, TU, and WIK background to three different wild-type strains: AB (Zebrafish International Resource Center), mixed background of AB/WIK/TU, or SAT (Zebrafish International Resource Center). Reference parameter values in Table 3 for 4, 7, 14 dpf fish were gathered using the AB strain fish.

*Drosophila melanogaster* (*w*^1118^) were raised at 23°on standard cornmeal-agar food under a 12/12 light/dark cycle.

*Caenorhabditis elegans* (*C. elegans*) were grown at 20°on nematode growth medium agar plates seeded with *Escherichia coli* OP50 as previously described ^129^.

### METHOD DETAILS

#### Behavior experiment

Larvae at desired age (4, 7, or 14 dpf) were transferred from petri dishes to behavior chambers at densities of 5-8 per standard chamber and 2-3 per narrow chamber with 25-30/10-15 ml of E3, respectively. After 24 h, behavior recording was paused for 30-60 minutes for feeding where 1-2 ml of rotifer culture was added to each chamber. Larvae were removed from the apparatus 48 h after the start of the recording.

Behavior measurement in this manuscript were collected from 27 clutches of zebrafish larvae between 7 to 9 dpf under constant darkness. 4 dpf and 14 dpf reference parameter values in Table 3 were collected from 10 clutches of zebrafish larvae under constant darkness. Finless data was generated using 4 clutches of larvae under constant darkness. For all experiments, a single clutch of larvae produces one experimental repeat with at least 3 behavior boxes each containing 5-8 larvae per standard chamber or 2-3 fish per narrow chamber.

For *Drosophila* recording, four flies were transfered to a narrow chamber. A small piece of water-dampened kimwipe was put at the bottom of the chamber to maintain humidity. A n acrylic plug was secured at the top to prevent them from escaping the chamber. We secured the chamber with the flies in the SAMPL apparatus and performed the standard SAMPL experiment using recording parameters provided in Table 2.

To image swimming *C. elegans*, eight starved N2 adult hermaphrodites were transferred to a narrow chamber filled with 15 ml M9 buffer (3 g/l KH_2_PO_4_; 6 g/l Na_2_HPO_4_; 0.5 g/l NaCl; 1 g/l NH_4_Cl) which was secured in the SAMPL apparatus as described above. Behavior recording was started immediately afterwards. Refer to Table 2 for SAMPL thresholds for *C. elegans* detection.

#### Fin amputation

6 dpf zebrafish larvae were anesthetized in 0.02% tricaine methanesulfonate (Syndel) and transferred to 3% Methylcellulose (Sigma). Fin amputation was done by removing pectoral fins using fine forceps (FST). Specifically, one pair of forceps was used to stabilize the head of the fish and a second pair was used to grab the joint and pull off the fins. Finless larvae were washed three-times in E3 and fed with cultured rotifers before behavior assessment at 7 dpf.

#### Video acquisition

Movie 1 was captured using Sigma fp digital camera (Sigma Co.). Video footage was edited and annotated using Premiere Pro (Adobe). Movies 2 & 3 was captured with the innate video capture function in SAMPL software using recording parameters described in Table 2. Movie 3 was edited using Adobe Premiere Pro (Adobe) to combine with timeseries data.

### QUANTIFICATION AND STATISTICAL ANALYSIS

#### Behavior analysis

Behavior data was analyzed using the Python analysis pipeline SAMPL_analysis_visualization. SAMPL_analysis() function was used to calculate swim parameters, extract bouts and inter-bout intervals (IBIs) from the raw data, and align swim bouts by the time of the peak speed.

Each run of the experiment (recording from “start” to “stop”) generates one data file (∗.dlm) containing recorded raw parameters including time stamp, fish body coordinates, fish head co- ordinates, pitch angle, epoch number and fish length at every time point. An epoch is defined by a duration where the number of detected pixels falls within the lower and upper threshold for recording, indicating detection of fish in the field of view.

To extract bouts from the raw data, first, swim features, such as speed, distance, trajectory, angular velocity, etc., were calculated using basic parameters and time interval. Next, epochs that were longer than 2.5 s, contain maximum swim speed greater than 5 mm/s, and pass various quality-control filters were selected for bout extraction. Epochs containing multiple bouts were segmented and truncated so that each detected bout contains data from 500 ms before to 300 ms after the time of the peak speed. Then, bouts containing 800 ms of swim data were aligned by the time of the peak speed and saved for further analysis.

All further quantification was performed on data during zeitgeber day, namely the 14 h light time for fish raising under 14/10 h light/dark cycle.

To calculate IBIs, epochs with multiple bouts are selected and the duration of swim speed below the 5 mm/s threshold between two consecutive bouts is calculated. A 100 ms buffer win dow is then deducted from each end of the duration to account for errors of swim detection (Figure 3A). Pitch angles during each IBI were averaged to generate an IBI pitch (Figure 3B).

Definition of other bout parameters can be found in Table 3. All bout parameters (except for kinetic parameters explained in the next section) reported in the main text and Table 3 are mean values across swim bouts collected from multiple experimental repeats. One experimental repeat is defined as behavior data collected from one clutch of fish over 48 h using at least three boxes.

#### Computation of kinetic parameters

To calculate larvae sensitivity to pitch changes (Figure 3), we plotted bout frequency as a function of IBI pitch. The data was modeled using a quadratic polynomial regression (least squares) defined by function:

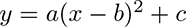

where the coefficient of the quadratic term *a* indicates sensitivity and the y-intersect *c* represents baseline bout rate.

To calculate steering gain (Figure 4), we first computed bout trajectory defined by the tangential angle of instantaneous trajectory. Pitch angles at time of peak speed were then plotted as a function of bout trajectories and modeled with linear regression (least squares). The slope of the best fitted line was termed the “steering gain.”

Time of peak angular velocity in Figure 5 was computed using adjusted angular velocity. First, pitch angles for each bout were smoothed by a window of 11 frames and used for calculate angular velocity. Next, we flipped the signs of angular velocity for bouts that started with nose-down rotation so that all bouts started with positive angular velocity. To calculate time of peak angular velocity, we took the median angular velocity at every time point across all bouts from the same experimental repeat and found the time for the peak. Peak angular velocity times across all experimental repeats were then averaged to generate mean peak time.

For fin-body coordination analysis (Figure 5), we selected swim bout that are faster than or equal to 7 mm/s. Bouts with steering rotations (posture change from −250 ms to 0 ms) greater than the 50^th^ percentile while having a negative attack angle were further excluded from analysis. To calculate fin-body ratio, we plotted attack angles as a function of early rotation. Attack angle is defined as the difference between bout trajectory and pitch at time of peak speed. Body change related to steering were calculated by subtracting pitch angles at time of max angular velocity by initial pitch. Attack angle-rotation plot was then fitted with a logistic function defined by

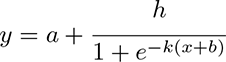

where *h* is the height of the sigmoid. Fin-body ratio was defined by the maximal slope estimated using *kh/*4.

To calculate righting gain and set point (Figure 6), righting rotation, defined by the pitch changes from time of peak speed to 100 ms after peak speed, was plotted as a function of initial posture. Righting gain was determined by the absolute value of the slope of the best fitted line. The x intersect of the fitted line determines the set point (Figure 6E, blue cross) indicating posture at which results in no righting rotation.

#### Estimating effects of sample size on statistical modeling of bout kinetics

For statistical analysis of swim kinetics (Figure 8A), the 7 dpf constant dark behavior dataset was sampled for 20 times at given sample number for calculation of swim kinetics and CI width. Specifically, sensitivity is determined by the coefficient of the quadratic term of the fitted bout-timing parabola as stated above. To plot estimated error as a function of the number of IBI, sets of data with N number of IBIs were sampled from the 7 dpf constant dark behavior dataset. However, different from the calculation of R^2^ above, the total dataset was sampled for 20 times for each desired number of IBIs (N). Regression analysis was performed on each set of sampled data to calculate sensitivity and its standard error. Estimated errors were used to calculate CI width at 0.95 significance level using normal distribution for each sampled dataset.

Similarly, steering gain and righting gain and their estimated errors were calculated from N number of bouts sampled from the original dataset. Estimated error was used to calculate CI width at 0.95 significance level for each sampled dataset. Sampling at each N was repeated for 20 times to generate error bars on the CI widths.

Fin-body ratio was calculated from N number of bouts sampled from the original dataset and repeated 20 times for each N. Because fin-body ratio is determined as the maximal slope of the sigmoid which is given by *kh/*4, the variance of fin-body ratio (slope) is calculated using formulation

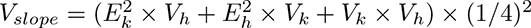

where *E_k_* and *E_h_* are the mean of *k* and *h* with *V_k_* and *V_h_*being their respective variance. Next, the standard errors of the fin-body ratio were calculated and used to estimate CI widths at 0.95 significance level.

To estimate effect sizes at given percentage of change (Figure 8B), an artificial data set was generated by altering the coefficient of interest while maintaining other coefficient as well as y residuals at given x values. N data points were drawn with replacement from each data set for calculation of kinematic parameters, which was repeated 200 times to generate distributions of parameters of interest. Effect sizes were determined using Cohen’s d:

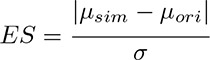

where *µ_sim_*and *µ_ori_*are the mean of parameter values calculated from respective data sets and *σ* is the standard deviation of all 400 calculated parameters. The whole process was repeated for 20 times to estimate the mean effect size at given sample size (N) and percentage of change. To reduce program execution time, we used a fixed 40 ms before time of peak speed as the time of max angular velocity for fin-body ratio calculation. Other kinematic parameters were calculated as described above.

## APPENDIX 1: HARDWARE DESIGN PRINCIPLES

### Camera

At the time of writing, the best price/performance ratio when using infrared light are the Sony Exmor line of complementary metal-oxide-semiconductor (CMOS) sensors. Sensors in the Exmor line are usually released as pairs, with a low-cost low-speed version of the same sensor available at the same time as a more expensive high-speed version. Our initial designed used the lower-cost IMX249 sensor; we have since switched to the faster IMX174 variant. These two sensors have a particularly large pixel size (5.86um), low noise (7e-), and a large well depth (32,513e-) allowing for exceptional dynamic range (73dB) and signal-to-noise ratio (45dB) at high-definition resolution (1936x1216 pixels). Quantum efficiency >900nm (i.e. the infrared range we will use) is 10%. Sony has released new sensors in the Exmor line regularly, but the trend has been to release sensors with increasingly small pixels. Thus for our purposes, the performance of the IMX174 remains unmatched.

Machine vision cameras are available with different interfaces used to stream data to a computer. The major difference between interfaces is the bandwidth available to each. The two most common interfaces for machine vision cameras at the time of writing are Gigabit Ethernet (125MB/sec) and USB3.0 (500MB/sec after overhead). Currently, there are commercially-available cameras with higher bandwidth interfaces utilize 10-tap CameraLink (850MB/sec), 10 Gigabit Ethernet (1250 MB/sec), 4xCoaXPress 2.0 (6,250MB/sec), and PCIe x8 (7,000MB/sec). Running our preferred IMX174 sensor at full resolution and speed for 8-bit images only requires 380MB/sec. Thus, USB3.0’s low cost and relative ubiquity made it the most attractive option for our apparatus.

There are a number of manufacturers that make cameras built around the IMX174 with a USB3.0 interface. Cameras from major manufacturers all conform to the GenICam standard making them largely interchangeable, particularly when using the Vision Acquisition software from National Instruments. We have successfully used cameras from Ximea (MC023MG-SY), Basler (acA1920-155um), and FLIR (GS3-U3-23S6M-C), others include SYS- Vistek (exo174CU3) and Daheng Imaging (MER2-230-168U3M). We have also used cameras ordered directly from different manufacturers – at a substantial discount – available via alibaba.com: Hangzhou Huicui Intelligent Technology Co. Ltd. (A7200MU130), Hangzhou Contrastech Co. Ltd. (Mars2300S-160um), Shenzhen Hifly Technology Co. Ltd. (MV-AU231GM). When ordering directly from manufacturers we specify Delivery At Place (DAP) shipping. The primary differences that we’ve encountered are whether a particular model implements binning or other on-camera computations, heat management, and different manufacturer-provided APIs. When we use multiple cameras from the same manufacturer on the same computer, we have also noticed that certain cameras will throw timeout errors on some USB ports but not others; shuffling cameras and ports has worked to solve this problem. At the time of writing, supply chain issues mean that most major camera companies quote long lead times, but cameras ordered directly through alibaba.com all shipped within two weeks.

### Illumination

Image quality is proportional to available light. Further, the size of the illuminated area defines the size of the field that can be imaged. Finally it is imperative for our experiments that from the fish’s perspective that the “dark” period is completely dark. We therefore chose 940nm LEDs as our source of infrared illumination. This left us with three options to build our illumination source: LEDs mounted on adhesive strips, “star” style LEDs with 1-4 dies on a single PCB, and a high-power LED array. The LED strips had too little illuminance for our purposes due primarily to the spacing of the LEDs. The high-power LED array had ample illuminance but generated so much heat that it required active cooling.

We developed a simple illumination module to provide diffuse IR light across a 50mm circle An LED mounted on a “star” PCB (Opulent LST-01F09-IR04-00, Mouser) provided ample light. We mount each “star” LED with thermal adhesive to a small heat sink (Ohmite SV-LED-325E) which in turn is glued to a Thorlabs adapter (SM1A6FW) to allows the wires to exit and the LED/heatsink to connect to collimation and diffusion optics. The heat sink is machined (either with a Dremel hand-held tool or a mill) on one side to allow the wires that power the LED to lie flat against the heatsink. We power multiple illumination modules in series using a constant current LED driver (LuxDrive BuckBlock 1000mA). Our illumination setup generates negligible heat and our modules run continuously for years.

Our imaging parameters are fixed across experiments and optimized to give the highest quality data we can achieve with our hardware. The gain of the camera is set either to its lowest value or just above to minimize noise. Our exposure time is either 750 µsec or 1msec, allowing for a crisp image in the face of the fastest movements that fish can make. The illuminated area is circular, but the image sensor size is rectangular. We therefore crop the sides of the image to produce a square that fits within the illuminated area.

### Lens

Our choice of lens was guided by the need to balance different demands:

1. wThe longer the working distance, the greater the space needed between the sample and the lens. We wanted our apparatus to fit length-wise on a 24 inch shelf, and so we needed to minimize the working distance.
2. The entire depth of the tank needs to be in focus, but not beyond that because we’d like to blur our LED.
3. The lens should be coated to pass IR light
4. The lens should be easy to mount to the base of the apparatus; mounting the lens instead of the camera allows drop-in replacement of cameras from different manufacturers, which have different positions of the tripod mount relative to the sensor.
5. The lens should have a simple way to mount an IR-pass filter (e.g. common thread).

Unfortunately, we were not able to find a single lens that met all of these criteria. Instead, we adapted a 50mm (Edmund Optics 67717) lens by placing a small Thorlabs tube (ThorLabs SM1-L03) between the lens and the camera. We mounted a 25mm IR pass filter (ThorLabs FGL830) inside the Thorlabs tube. By moving the lens farther from the sensor we decreased the minimum working distance sufficiently. Finally, the Thorlabs tube allows us to mount the lens to the breadboard directly.

### Behavioral arena

To maximize the amount of time the fish swam in a plane orthogonal to the camera, we used rectangular chambers. Initially we chose glass colorimeter cuvettes (Starna Cells Inc, Atascadero CA): they are made of an inert material (glass) and come in a variety of sizes. Due to supply chain issues, we switched to custom-fabricated chambers, plans attached. We now assemble these from laser-cut acrylic, cementing a front and back side to a u-shaped piece that forms the other sides. These chambers are considerably cheaper and less prone to breakage than glass and can be rapidly modified to allow for different experiments.

### Enclosure

We designed a custom aluminum base with tapped holes for post-holders for the IR LED, chamber holder, and camera/lens/filter holder. We used custom-cut extruded aluminum rails to frame the sides and top. The sides are made of black foam-core sized to fit in grooves in the breadboard and rails. The top rails have a cross piece that holds the LED strip used to provide circadian lighting. All parts are fabricated to order by Base Lab Tools Inc (Stroudsburg PA). The top is a steel tray fabricated to order by MetalsCut4U (Avon Lake OH). Our current enclosure took roughly three months to prototype before settling on the final design.

### Shelving and fleet organization

We have organized our fleet of apparatus to sit on mobile wire shelving. Currently, we use 36”x24”x81.5” adjustable wire shelving units (McMaster Carr, Robbinsville NJ). We prefer to have the shelving on casters as it makes accessing the back of the units considerably easier. Shelving is organized such that one computer and three apparatus sit on a single shelf. Enclosures on a given shelf are color-coded (blue, gold, and red) so that each apparatus can be uniquely identified by a color/shelf/module combination; this also facilitates wire labeling. Each shelf has its own power strip that controls the computer, the IR lights, and the white LEDs; all strips plug into a single uninterruptible power supply (APC SmartUPS 1000C).

Our aim in specifying module size was to ensure that multiple investigators could set up experiments simultaneously, and to minimize the cost One unit has four shelves so that a single “module” consists of four computers and twelve apparatus. Each module has a dedicated monitor/keyboard/mouse on an adjacent desk, shared by the four computers using a KVM switch (IOGEAR GCS1794). A module has its own dedicated unmanaged Ethernet switch (NETGEAR GS110MX) that allows Gigabit speed communication between computers and 10 Gigabit speed between modules.

### Computer hardware

Computer hardware was chosen to ensure adequate performance while minimizing cost, noise, and size. We found that building our own computers was the only path forward in the face of supply chain issues and strict optimization criteria. We opted to build around what was, at the time of writing, the previous generation of AMD microprocessors (Ryzen 7 5700G) cooled by a Noctua NH-L9a-AM4 fan (to minimize acoustic noise). We chose a Mini-ITX form factor motherboard that allowed us to use a small case (Cooler Master NR200). Other parts (64GB RAM, SSD, power supply) were chosen based on availability; a full parts list is attached (Table 1). We recommend using https://www.pcpartpicker.com to minimize cost and ensure compatibility of different components. All computers run Windows 10 Professional (Microsoft, Redmond WA).

## APPENDIX 2: ACQUISITION SOFTWARE DESIGN PRINCIPLES

### What we don’t measure

To extract the maximum amount of useful information about posture and locomotion with the minimum amount of overhead we had to be selective about what we measure. Our imaging field is located in the center of the arena; fish that swim at the bottom, top, or sides of the tank where there is a boundary are excluded from tracking. While multiple fish swim in the same arena, we do not take data when more than one fish is in the imaging field to sidestep the need to track fish identity. Our arena is sized to allow fish to swim freely but its shape (a rectangular solid) encourages fish to swim in line with the imaging plane; we exclude frames where fish turn away from the field of view (i.e. are swimming towards/away from the camera). Finally, capturing the full range of rapid propulsive undulations of the fish tail requires a frame rate of 500Hz-1kHz ^130, 131^. As changes to posture and locomotion are much slower, we opted to record at 160Hz. Together, these choices allowed us to optimize our algorithms to achieve the speed necessary to process video in real-time.

### Algorithms to measure posture and position

Our apparatus extracts the position and pitch orientation of zebrafish in real-time over days using a simple set of common machine vision processing steps:

1. Measure the absolute difference between the current frame and the background (fish-free) image.
2. Threshold the difference image such that all small differences are set to zero.
3. Dilate the image three times in succession to remove any larger clumps that are still smaller than a fish.
4. Extract and quantify all particles in the image.

Real-time video processing allows efficient data extraction during video acquisition. Our design of the architecture is further discussed in the section: Optimizations for speed.

Below we detail a number of additional processing and optimization steps to ensure that we maximize useful data.

### Measuring the pitch of the fish

To extract the pitch (the angle of the fish with respect to the horizon), we perform the following steps to ensure that the sign and magnitude of the angle is correctly assigned:

1. Fit the particle with an ellipse and extract the angle of the long axis with respect to the horizon.
2. Threshold the original difference image again to identify the pixels that correspond to the head of the fish.
3. Using the head and body (X,Y) coordinates determine whether the fish was facing to the left or right.
4. Assign the correct angle and sign such that nose-up posture is always positive and nose-down is always negative.

These steps ensure that the data saved follows a simple and intuitive convention for posture.

### Optimizations for speed

To optimize our code for speed, we use a set of thresholds to rapidly evaluate and reject frames

1. Before any processing, we sum the pixel values in the frame. If it is too low (no fish in frame) or too high (more than one fish in the frame) we reject the frame.
2. After the particles are identified we reject the frame if a particle is touching the edge (fish partially out of frame), if there is more than one particle (multiple fish) or if the length of the particle is too short (fish bending in/out of the field of view). We define an epoch as a set of continuous frames that pass all our exclusion criteria (i.e. that contain a single fish in frame). Epoch duration is tracked and, when too short, can be rejected.

In addition to optimizing the algorithm, we adopted a producer-consumer architecture to decouple video acquisition from video processing and saving data. Our software runs two routines: the “producer,” which acquires frames from the camera and places them in a queue in memory, and the “consumer” that extracts each frame from the queue and processes it in turn. Our program monitors the size of the consumer buffer and, if it has less that 10% free, pauses the producer routine for 15 seconds to allow the buffer to clear. In this configuration, the performance ceiling shifts from CPU speed (i.e. how quickly can a frame be processed) to the amount of RAM available (i.e. how many frames can be queued). At the time of writing this, doubling the amount of RAM is considerably less expensive than doubling CPU performance. The choice of architecture thus brings down the cost of the computer.

### Saving raw video

While the bulk of our experiments rely on real-time processing of video it is often useful to save the actual data. Further, we wanted to be able to set user-defined criteria to determine in real-time which videos were worth saving. Leveraging the producer-consumer architecture, our software contains a routine that independently buffers the frames being analyzed and, if, the video to be saved meets user-defined criteria, will pass the frames to an independent program to write them to disk. For example, we can ensure that the video to be saved is of a certain length. Similarly, we can filter the video images ^132^ to determine if the target is in crisp focus (useful for larger arenas, or higher magnification) and only save high quality videos. By separating video writing from acquisition and processing, comparatively slow operations such as video compression and/or saving video to a network-accessible shared drive do not compromise performance.

### Apparatus control software

Our algorithm relies on common and mature image processing routines and could be instan- tiated in any modern programming language. Since we had run this algorithm for the better part of a decade we were confident that it was sufficiently stable to compile into a distributed executable, which would greatly simplify deployment to a fleet of apparatus. Our original im- plementation was written in LabVIEW (National Instruments, Austin TX) which was stable and accommodated all the lab’s hardware changes for the past decade. We therefore opted to up- date the LabVIEW code, which we distribute both as source and executable versions. Running the executable requires each computer to have the LabVIEW Runtime Engine (free download) installed, as well as a license for NI Vision Acquisition software (NI 778413-35) and the Vision Development Module Run-Time engine (NI 778044-35).

### User interface

We designed the interface to enable easy initialization of experiments, rapid graphical and quantitative visualization of video processing and performance, and to minimize error. Launching the executable starts the program, which allows the user to fill out various text, numeric, and drop-down fields that describe the experiment. The user then monitors the video feed until no fish are in frame and then selects that image as the background. We have found that this initial bit of monitoring both compensates for slight day-to-day differences in arena placement. More importantly, it forces the user to monitor the live feed at the beginning of each experiment, a useful bit of mindfulness that minimizes lost data. Once running, the user can: monitor the output of each step in the processing algorithm graphically, monitor the number of times the consumer buffer has overflowed (usually zero), update the text fields, and stop the program. Hardware parameters are stored in a text file that can be easily edited directly. Experiment parameters are similarly saved to text files and can be reloaded to save time.

We have implemented a number of user interface items to minimize confusion in the face of a fleet of instruments. First, we have color-coded versions of the executable (blue, gold, and red) where the background clearly differentiates the version. Each version has its own configuration file that, during setup, is coded to a particular apparatus. Thus the user is always aware of which apparatus they are interfacing with based on color cues. Next, we added a “debug” button to the front panel that allows for direct monitoring and editing of all program variables. In “debug” mode the user has the option to save raw video.

## APPENDIX 3: HARDWARE ASSEMBLY GUIDE

In this Appendix, we walk through box assembly and recording settings. Refer to Appendix 5 for executing experiments and SAMPL data analysis.

### Hardware assembly

Our design of hardware allows connecting up to three SAMPL boxes to one computer while using one set of power supplies (for IR LED and daylight LED). See Movie 1 for video instruction on box assembly.

1. Camera module

a. Attach 1x 1.5 inch post (TR1.5) to the camera module holder (SM1RC). Screw in tightly.
b. Assemble lens and camber. Sequentially connect parts below: camera lens, SM1A10 adapter, IR filter, SM1-L03 extension tube, assembled camera module holder, SM1A9 adapter, and the camera.
2. IR illumination module

a. Attach wired IR LED to the heatsink and SM1A6FW adaptor (see below for instruction).
b. Carefully mount the condenser into SM2L05 tube.
c. Assemble the IR module by sequentially connecting parts below: IR kit, SM1M10 tube, SM1A2 adapter, SM2L20 tube, and the mounted condenser.
d. Tightly attach TR1 post to SM2RC holder.
3. Chamber holders

a. Take off rubber covers on the tip of the screws on the chamber holders (FP01).
b. Mount holders onto TR1 posts using 8-32 screws.
c. Assemble the IR module by sequentially connecting parts below: IR kit, SM1M10 tube, SM1A2 adapter, SM2L20 tube, and the mounted condenser.
d. Tightly attach TR1 post to SM2RC holder.
4. Put together the box

a. Mount one of the short rails between two long rails using right angle brackets and T-nuts.
b. Mount the other two short rails onto the long rails using slotted cubes and low-profile cap screws (SH25LP38).
c. Adjust the position of the middle rail so that is approximately 13 cm away from one end of the frame.
d. Attach post holders to the base plate using cap screws (SH25S038). Note that the one for the camera module is the longer post holder (PH1.5).
e. Mount 4 medium rails onto the base plate using standard cap screws.
f. Insert all the modules onto the base plate. Connect USB cable to the camera.
g. Make a notch in the middle of the shorter side of a small panel and insert it between the rails on the side of the camera module.
h. Insert 2 large side panels.
i. Attach daylight LED to the top frame (see below for instruction).
j. Pass IR and daylight LED wires through the front notch of the baseplate.
k. Insert the front panel.
l. Attach the top frame.

### IR light wiring

Solder 2x 9” wires onto the IR LED “star.” Attach IR LEDs to heatsinks using HexaTherm tape. Note that in order to pass the wires through the heatsinks and the SM1A6FW adapter on the opposite end, the ears of the Ohmite heatsink need to be trimmed down a little. When done, attach the heatsink to the adapter using thermal epoxy. To simplify light wiring, we use one 1000 mA BuckBlock to dirve 3 IR lights in series for 3 boxes on the same level of the shelf. To do this, one needs 2x 7” wires to connect adjacent IR cables and 1x 22” wire connecting the further IR to the BuckBlock. Use another 8” wire to connect the closest IR to the BuckBlock. We recommend using XT60H connectos to link these wires to the IR light cables and connect wires to the BuckBlock for the ease of troubleshooting and replacement. Finally, connect the BuckBlock to 12 V 2 A power supply through pigtail adaptors.

### Daylight wiring

Each box uses a strip of 6 daylight LEDs. Our choice of daylight LED comes with double sided tape already attached to the back side of the LED which is used to install LED strips to the top frame. To wire daylight LED strips, solder 2x 20” wires to the LED strip. Heat shrink sleeves can be used here to strengthen connections. Twist the wires at the end close to the LED. This helps with cable management in the box. Bend the cables 90 degrees in the XY plane (perpendicular to the illumination direction) so that the wires won’t get into the field of view.

To simplify light wiring, we use one 12V 1A power supply to drive 3 LED strips in parallel. To do this, cut 1x 27” wire for connecting the positive end of the DC plug to the LED strip. Prepare 3 wires for the negative end of the strips each measured 10”, 18”, and 27”. Insert one end of all three wires for the negative end into the negative terminal of a pigtail connector, connect the other ends to the LED cables. For positive end, we recommend using T tap connectors (B085XGYW1B) which allows easy disconnection.

### PC setup

Assemble computer parts. Make sure 1 PCI-e USB card is installed into each PC. Connect power cable and Ethernet cable. If desired, connect 3 cameras to three different USB BUS on the PC: specifically, one to a PCI-e USB card, one to a USB 3.0/3.1 port on the motherboard in the back of the PC, and one to a USB 3.0 port on the front panel. If desired, connect to the KVM switch.

Turn on the PC, setup Windows. If necessary, change settings below to achieve peak performance: select AMD High Performance in Power Settings; set Sleep time to Never; set hard disk sleep time to 0 in Advanced Power Settings.

### Install behavior programs

We provide three executable programs (Blue, Gold, Red) that can run simultaneously on the same PC. Refer to the Key Resources Table for access to the programs. To install executables, download *.exe files and corresponding configuration files (* Configuration.ini). Create a folder under C:/ and move configuration files to C:/Data/. Install required NI software and activate: LabVIEW Runtime, Vision Acquisition, and Vision Runtime. Restart computer.

Open NI Max, rename cameras to camBlue, camGold, and camRed. Set camera settings:

- Field of view - X: left = 360; resolution = 1216
- Field of view - Y: top = 0; resolution = 1200
- Under Acquisition attributes - Receive time stamp mode = System time
- Under Camera attributes - Analog control - Gain = 1; Black level = 1 (if applicable)
- Under Acquisition Control - Exposure time = 1000; Trigger activation = Rising edge; Frame Rate = Freerun (for 166 Hz with our cameras of choice, or set to desired frame rate)

Open configuration files and set box number to desired values. We use box number as a unique identifier for different behavior boxes. Check camera name to make sure it’s the same as the corresponding camera names in NI Max.

Open behavior programs, now that you should see images showing up on the preview windows.

### Camera calibration

Once the apparatus has been assembled and software has been installed, align the field of view (FOV) to the center of the IR light circle. Raise or lower the post holding the camera module to center the FOV in Y and roll the module to level it.

Next, calibrate the scale of the FOV to 60 pixels/mm. To do this, secure a micrometer in a chamber and place it into the box. Snap a picture of it using NI Max, then measure the scale using the image of the micrometer. If necessary, loosen the SM2RC adapter and move the camera and lens forward or backward to achieve the correct scale.

Illumination adjustments should be completed with the behavioral arena in place. To calibrate exposure, first ensure the correct IR light is in use and set the aperture ring between f/16. In NI Max, the peak of the image histogram peak should be around 128 (the middle of the 8-bit range). If necessary, exposure can be reduced by lowering exposure time or increased by open- ing up aperture to f/11.

### Network setup

We use a Synology data server as a repository to store behavior data. Hard drives are setup as RAID 10. Each SAMPL rack has its own ethernet switch, which can be connected to other switches as necessary.

## APPENDIX 4: DATA ANALYSIS SOFTWARE

In this appendix, we discuss algorithms for the data analysis and plotting software. We assume that the user is working with data from larval zebrafish here. If not, the specific parameters identified here are unlikely to translate as other organisms move differently but can nonetheless be used as a starting point. Refer to Appendix 5 for instruction for use. Refer to the Key Resources Table for access to the code.

### Read DLM files

Each SAMPL session (from Start experiment to Stop) generates one tab-delimited (i.e. .dlm) file. Each time point appears as a row of tab-separated values in the .dlm file. Columns, from left to right, are time stamp, fish number in the field of view (FOV), pitch angle (0-90°), x coordinate for body, y coordinate for body, x coordinate for fish head, y coordinate for fish head, raw fish angle (0-180°), epoch number, and estimated fish length.

Each .dlm’s data is loaded as a Pandas DataFrame for further analysis (see src/SAMPL_analysis/preprocessing/read_dlm.py for details). Each raw DataFrame contains multiple epochs. An epoch is defined as duration when a fish is detected in the FOV. See Appendix 1 for details on the algorithm for animal detection.

### Extract epochs

We calculated swim attributes, such as angular velocity, swim speed, instantaneous displacement, etc., from recorded pitch angles and fish body coordinates. To extract quality epochs from the recorded data, epochs are analyzed and passed through several quality control filters:

1. each epoch is truncated by 50 ms at both the start and the end to eliminate frames when fish is entering/exiting the FOV;
2. epochs with duration shorter than 2.5 s are excluded (for 1 & 2, see function raw_filter());
3. epochs with frame drop greater than 3 frames are excluded;
4. epochs with direction of fish translocation opposite to where the head points toward are dropped (for 3 & 4, see function dur_y_x_filter());
5. epochs with aberrant displacement jumps are excluded;
6. epochs with improbably large angular velocity greater than 250°/s or angular acceleration larger than 32000°/s^2^ are excluded (for 5 & 6, see function displ_dist_vel_filter()).

All the processes above can be found in the script src/SAMPL_analysis/preprocessing/analyze_dlm_v4.py.

### Get bout and inter-bout data

Epochs that pass the quality control are used to extract swim bouts using function grab_fish_angle() under src/SAMPL_analysis/bout_ analysis/grab_fish_angle_v4.py.

We use a swim speed threshold of 5 mm/s to determine swim windows. Adjacent swim windows with intervals smaller than 100 ms are combined. Next, we find the time of the peak speed for each swim window and extract frames in a range of 500 ms before to 300 ms after that. Inter-bout intervals (IBI) are determined as time between adjacent swim bouts with a 100 ms buffer window deducted from both the beginning and the end and IBI data is extracted accordingly. Baseline is considered the time during which larvae swim slower than 2 mm/s and baseline parameters are extracted accordingly.

Note that an epoch can only contain a single detected fish. The number of swim bouts extracted from an epoch various extensively depending on the quality of the epoch (and behavior of fish). Having too many fish in the chamber may lead to low yields of aligned bouts despite having a large number of epochs. For details of fish detection, refer to Appendix 1.

### Export analyzed results

Numerous attributes are saved as DataFrames under keys in HDF5 format files using our analysis pipeline. Once the analysis is complete, three output data files are generated: all_data.h5, bout_data.h5, and IEI_data.h5.

The all_data.h5 file contains epoch-based data including raw data from DLM files, epoch attributes, baseline angular velocity, etc. The bout_data.h5 file includes bout attributes and aligned bout data such as pitch angles and speed. The IEI_data.h5 file contains all inter-event interval (IEI) data, or IBI. Refer to docs/ for a complete list of saved attributes and their description. In addition, a metadata table including recording frame rate, number of aligned bouts, and other information is generated and saved to the same directory.

All results are saved as “long format” DataFrames with each row representing a time point or a bout/IEI, depending on the type of the result (one value per timepoint vs per bout/IBI). Values of multiple aligned bouts are stored in successive rows.

All functions above can be called with script src/SAMPL_analysis/SAMPL_analysis.py. Refer to Appendix 5 for running instructions. For a record of analyzed files, frame rate, number of aligned bouts, etc., refer to the log file generated under src/.

### Load analyzed data and calculate parameters

We include several plot functions under src/SAMPL_visualization/ that calculate and plot all the parameters we report in the main text. These functions require an input of a root directory containing analyzed data. For recommended behavior data structure, see Appendix 5.

Once data is found, plot functions get frame rate from metadata files and calculate the index of time of peak speed which is used to calculate the number of aligned frames and initialize other constants. Note that plot functions only read one frame rate for all the data to be plotted. Therefore, make sure all experiments are done at the same frame rate. To combine results from different frame rates for plotting, extract parameters of interest separately for experiments with different frame rates and concatenate the results afterwards. We only plot zeitgeber day data in this version of the code. Users may modify the day_night_split() function to extract zeitgeber night results if intended.

To load analyzed swim bouts and IBI, we loop through all subfolders under the root directory and read DataFrames from HDF5 files, extract and calculate desired parameters and concatenate results. Each plot function extracts parameters in different ways.

For time series values to be plotted as a function of time, data is loaded from the all_data.h5 file. The key prop_bout_aligned contains propulsive bouts that have been aligned and grabbed_all includes all epochs that contain swim bouts. See plot_timeseries.py for examples.

Bout parameters, such as speed, displacement, pitch angles and attack angles, are also extracted from prop_bout_aligned key containing aligned swim bouts. We use a dedicated function for calculating these swim parameters: extract_bout_features_v4(). These parameters can be further used to get steering and righting gains. See get_kinetics() for more. Note that some parameters are determined by specific time points (such as initial pitch, post-bout pitch, etc.). To determine frames that are the closest to these time points, we use half round up for rounding.

IBI data is loaded from the IEI_data.h5 file under key prop_bout_IEI2. For bout timing estimation, we calculate bout frequencies as reciprocals of bout intervals (IBIs). See plot_bout_timing.py and plot_IBIposture.py for examples.

To calculate fin-body coordination, users need to determine how the rotation is calculated. One way is to use rotation to time of peak angular velocity which requires estimation of time of peak angular velocity. To do this, we first calculate angular velocity using smoothed pitch angles and adjust the signs so that values are positive before time of the peak speed. Median of angular velocity time series from the same experimental repeat (see Appendix 5 for data organization) is used to find time of peak angular velocity. Lastly, we average results across experimental repeats to determine the peak angular velocity time. However, this calculation requires a large amount of bout data. Alternatively, one may use a fixed value for time of peak angular velocity. Generally, we found −50 ms (50 ms before time of peak speed) to be a good value to use. Once the time of peak angular velocity is determined, rotation is calculated by pitch change from 250 ms before peak speed to time of peak angular velocity. Some scripts have the option to sample data from each experimental repeats. See Appendix 5 for instruction.

### Visualize results

We use the Seaborn package for data visualization ^133^. Each plotting script generates a folder under figures/ and saves figures as PDFs. Below is a list of available plotting functions and their descriptions. For more details, refer to the README document.

1. plot_timeseries.py plots basic parameters as a function of time. Modify all_features to select parameters to plot. This script contains two functions: plot_aligned(), plot_raw(). Change variable all_features to select parameters to plot.
2. plot_parameters.py plots swim parameter distribution and 2D distribution of parameters for kinetics calculation. This script contains function: plot_parameters().
3. plot_IBIposture.py plots Inter Bout Interval (IBI; aka inter-event interval, IEI) posture distribution and standard deviation. This script contains function: plot_IBIposture(). This script looks for prop_Bout_IEI2 in the prop_bout_IEI_pitch data which includes mean of body angles during IBI. When input root directory contains multiple experimental repeats, the scripts allows sampling of IBIs from each repeat by specifying argument sample_bout.
4. plot_IBIposture.py plots Inter Bout Interval (IBI; aka inter-event interval, IEI) posture distribution and standard deviation. This script contains function: plot_IBIposture(). This script looks for prop_Bout_IEI2 in the prop_bout_IEI_pitch data which includes mean of body angles during IBI. When input root directory contains multiple experimental repeats, the scripts allows sampling of bouts from each repeat by specifying argument sample_bout.
5. plot_bout_timing.py Plots bout frequency as a function of IBI pitch and fitted coefficients of function. This script contains function: plot_bout_frequency(). When input root directory contains multiple experimental repeats, the scripts allows sampling of bouts from each repeat by specifying argument sample_bout.
6. plot_kinematics.py Plots righting gain, set point and steering gain. This script contains function: plot_kinetics(). When input root directory contains multiple experimental repeats, the scripts allows sampling of bouts from each repeat by specifying argument sample_bout.
7. plot_fin_body_coordination.py Plots attack angle as a function of rotation and calculates fin-body ratio. Rotation is calculated by pitch change from −250 ms to −40 ms. This script contains function: plot_fin_body_coordination(). For reliable sigmoid regression, 6000+ bouts is recommended. When input root directory contains multiple experimental repeats, the scripts allows sampling of bouts from each repeat by specifying argument sample_bout.
8. plot_fin_body_coordination_byAngvelMax.py Plots attack angle as a function of rotation and calculates fin-body ratio. Rotation is calculated by pitch change from −250 ms to time of max angular velocity. For reliable sigmoid regression, 6000+ bouts is recommended. When input root directory contains multiple experimental repeats, the scripts allows sampling of bouts from each repeat by specifying argument sample_bout.

## APPENDIX 5: STANDARD OPERATING PROCEDURE FOR RUNNING EXPERIMENTS AND ANALYZING DATA WITH SAMPL

In this appendix, we provide a step-by-step instruction for running experiments and analyzing SAMPL data. Refer to the Key Resources Table for access to SAMPL analysis and visualization scripts.

### Experimental design

SAMPL experiments usually involve comparing behaviors of two or more groups of fish with different mutations, transgenic backgrounds, or manipulation. We suggest first deciding *a priori* on the total number of bouts required to resolve differences Figure 8 with the desired power. Typically, one SAMPL experimental repeat containing two 24-hour sessions using 3 boxes with 5-7 larvae per box yields 3000-6000 bouts, which is usually sufficient for parameter calculation Figure 8. However, multiple factors can affect data size per repeat, such as: manipulations (mutation/drug treatment), the throughput of manipulation, the availability of apparatuses, and the number of larvae with desired background per clutch. We therefore suggest running a pilot experiment first to determine the number of bouts that can be expected per box. Once done, we suggest defining an “experiment” with respect to the desired number of bouts, which will specify the number of boxes and larvae per box required. Outlier boxes with too few or too many bouts (e.g. more/less than 2SD) can then be excluded from further analysis according to pre-determined criteria. Finally, we recommend running the full “experiment” multiple times to ensure that the findings are reproducible, and to report the variance across estimated parameters. Certain circumstances may be ill-suited to this approach: for example, if particular genotypes of larvae are especially rare, such as in the case of doubly biallelic mutants, or genotypes that simply swim drastically less. In such cases one can combine swim bouts across experimental repeats, and report the estimated error in pararmeter estimates using statistical resampling techniques such as the jackknife.

### Running an experiment

One typical SAMPL experimental repeat contains two 24-hour sessions. We suggest running zebrafish larvae at one of 3 time points: 4-6 dpf, 7-9 dpf, or 14-16 dpf. Larvae should be given 30 minutes of access to food before being placed into chambers. We suggest putting 5-8 larvae into one standard chamber and 1-3 larvae in one narrow chamber to maximize data yield. Behavior recording requires having a single fish in the FOV at a time. Appearance of additional larvae will disrupt fish detection. We suggest transferring 25-30/10-15 ml E3 medium into each standard/narrow chamber to account for evaporation and maximize likelihood of fish swimming in the FOV. Throughput of the apparatus can be found in Figure 2 (standard chamber based on 58 larvae; narrow chamber based on 23 larvae).

With SAMPL, one computer can control up to three behavioral apparatus, or “boxes.” Once the fish chamber is put into the box and secured, open the program (Blue, Gold, Red) corresponding to the box to run on the computer controlling the boxes. Enter experimental information in the window opened: Genotype (experimental conditions), Cross ID, Fish number, etc. Set the destination folder for data storage. Choose the desired Light-Dark (L/D) cycle from one of the followings: L/D, L/L, or D/D. Adjust daytime light connection/timer accordingly. Use fish size toggle to select thresholds for fish detection: use Small fish for larvae younger than 12 dpf and Big fish for those that are older. To start recording, click Select Background when there’s no fish in the FOV.

Larvae older than 5 dpf should be fed every 24 hours with 1-2 ml of diluted cultured rotifers. To feed fish, click Stop program to stop the current session. Feed with rotifers and allow a pause of 30 min before re-starting the experiment.

At the end of the experiment, click Stop program and remove fish from the box. Each session (from Start to Stop program) generates one .dlm data file and a corresponding .ini metadata file.

### Software requirement for data analysis

To analyze behavior data using code provided, one needs Python 3, analysis scripts, and various Python modules. An integrated development environment (IDE) is recommended to edit, debug, and run the code. If you don’t have a personal preference, we recommend using Visual Studio Code (Microsoft). Analysis and visualization code was developed using Python 3. For the ease of package management, we suggest the use of environment management tools, such as miniconda.

The most recent version of the code we use to analyze SAMPL data can be found online at https://. Download the entire directory by pressing the green Code button and downloading the ZIP file (orange box) so that you can make changes as needed for your project. The src folder contains all scripts. The sample figures folder contains examples of plots from the visualization functions. Please refer to the README for instructions and user guides.

To set up a virtual environment, open a new terminal or use the terminal in your IDE, and type:

conda create -n <MYENV>

where

<MYENV> is substituted with any desired name for the environment. Next, activate this environment

conda activate

<MYENV>

and install packages required for analysis and plotting using

conda install

<PACKAGE>

Below is a list of required packages ^134–140^ other than those included in Python 3.10.4:

- astropy=5.1
- pandas=1.4.4
- pytables=3.7.0
- matplotlib=3.5.2
- numpy=1.23.3
- scipy=1.9.1
- seaborn=0.12.0
- tqdm=4.64.1
- scikit-learn=1.1.1

For a complete list of packages, refer to the environment.yml file.

### Bout analysis

Analysis and plotting scripts support two types of data structures. The first option is one root directory containing all data files:

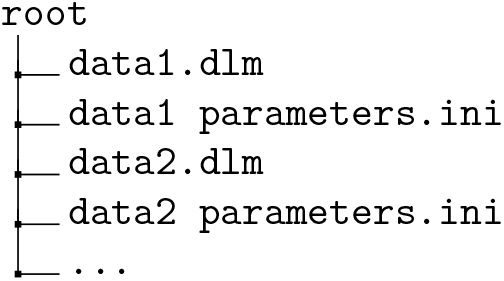

The second is a root directory containing subfolders with the necessary files indicating experimental repeats:

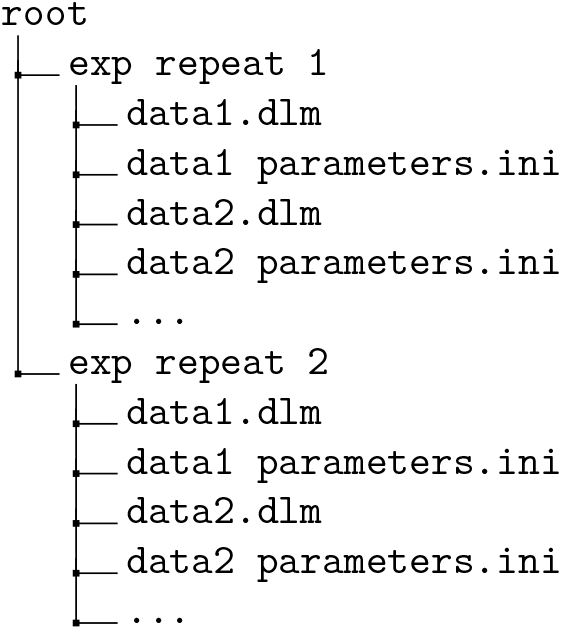

Run the analysis script …/src/SAMPL_analysis/SAMPL_analysis.py and input data directory (directory of the root folder) and the frame rate as instructed. This function aligns bouts in .dlm files within a directory so that peak speed is at time 0 ms, with 500 ms of activity before and 300ms of activity after. It is important to note that all files in the same subfolders under the input directory will be combined to extract bout parameters. The analysis script will take the submitted directory and analyze all data files within it, including all subfolders in its search, regardless of depth. Subfolders can be used to separate analyses, experimental conditions, or repeats. Data with different frame rates should be analyzed separately to ensure proper parameter calculation, as only one can be used at a time.

The program will skip the current .dlm file if it fails to detect a bout in it. However, errors are expected if files contain too little recorded data to extract a bout. Therefore, we suggest removing any .dlm files that are smaller than 1 MB.

When analysis is done, it will save three data files (.h5), four catalog files (.csv), and two metadata files (.csv) under the same directory as the data is in. Below is an example of an analyzed directory:

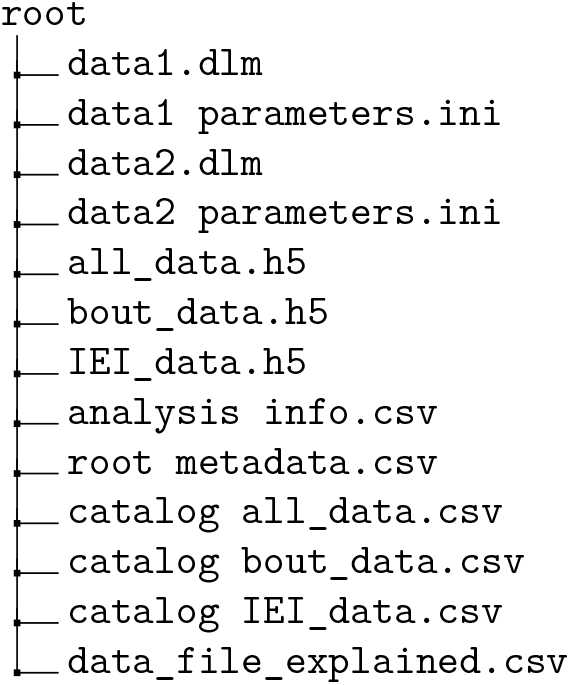

### Visualizing results

After analysis, the scripts under the visualization folder are used to extract swim parameters and kinetics, and visualize them. For more detailes, refer to Appendix 4 and the README document. Each function can be run individually and will ask for the directory path to your data (see the Bout analysis section above). Alternatively, use plot_all.py to plot all figures.

If the data size from a single repeat is not adequate for parameter calculation, we suggest combining data from multiple repeats and use sampling techniques such as Jackknifing for error estimation.

---

**Figure S1:**
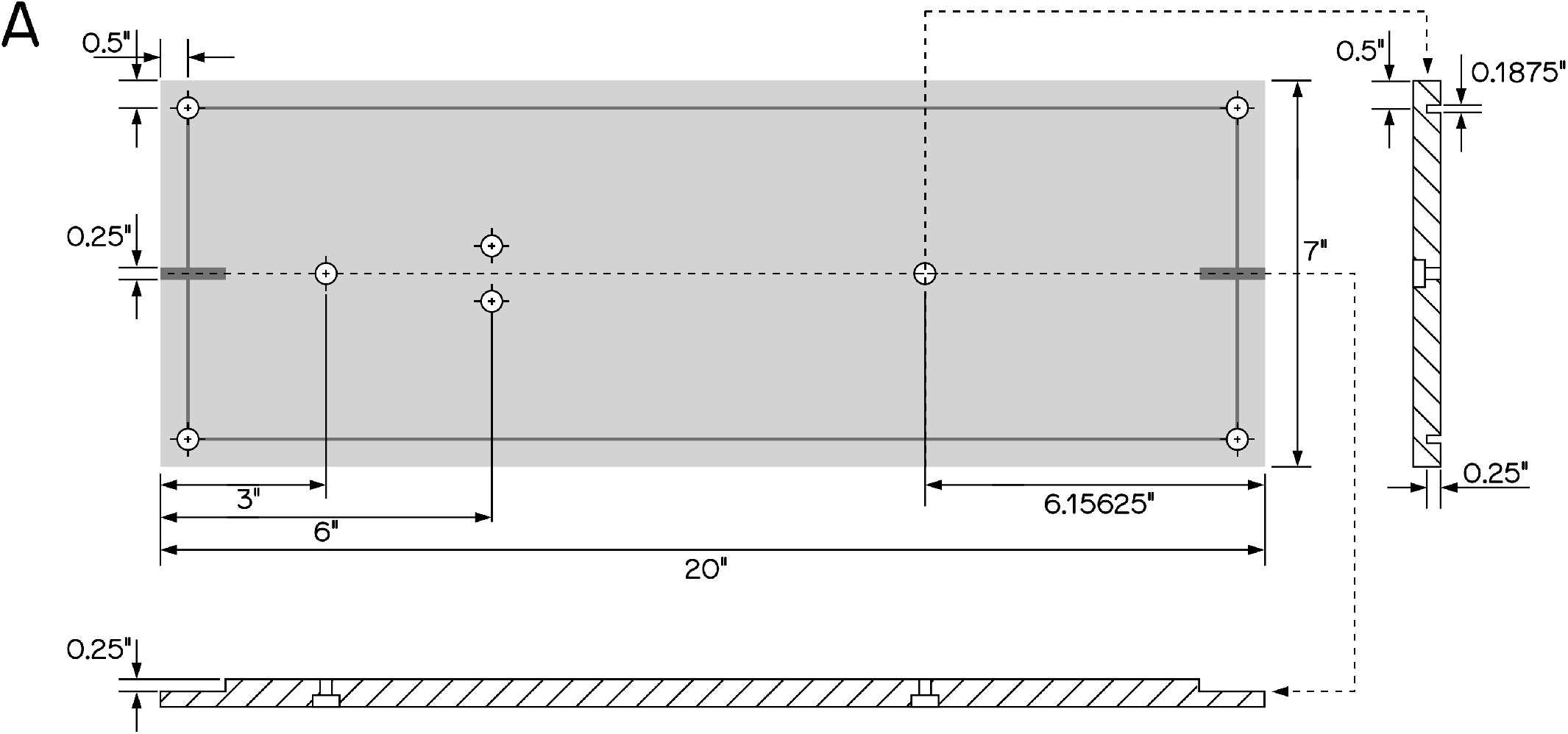
Custom breadboard for SAMPL base. (A) Custom aluminum breadboard, not anodized, 0.5” thick. All holes (8 total) counterbored for 1/4”-20 cap screw. Grooves to be cut on the side of the breadboard OPPOSITE to the counterbore.

